# Consequences of producing DNA gyrase from a synthetic *gyrBA* operon in *Salmonella enterica* serovar Typhimurium

**DOI:** 10.1101/2020.12.08.416404

**Authors:** German Pozdeev, Aalap Mogre, Charles J Dorman

**Affiliations:** Department of Microbiology, Moyne Institute of Preventive Medicine, Trinity College Dublin, Dublin 2, Ireland

## Abstract

DNA gyrase is an essential type II topoisomerase that is composed of two subunits, GyrA and GyrB and has an A_2_B_2_ structure. Although both subunits are required in equal proportions to form DNA gyrase, the *gyrA* and *gyrB* genes that encode them in *Salmonella* (and in many other bacteria) are at widely separated locations on the chromosome, are under separate transcriptional control and are present in different copy numbers in rapidly growing bacteria (*gyrA* is near the terminus of chromosome replication while *gyrB* is near the origin). We generated a synthetic *gyrBA* operon at the *oriC*-proximal location of *gyrB* to test the significance of the gyrase gene position for *Salmonella* physiology. Producing gyrase from an operon did not alter growth kinetics, cell morphology, competitive fitness index, or sensitivity to some gyrase-inhibiting antibiotics. However, the operon strain had altered DNA supercoiling set points, its SPI-2 virulence genes were expressed at a reduced level and its survival was reduced in macrophage. The *gyrB* gene could not be deleted from its *oriC*-proximal location, even in a *gyrB* merodiploid strain. We discuss the physiological significance of the different *gyrA* and *gyrB* gene arrangements found naturally in *Salmonella* and other bacteria.

## Introduction

DNA gyrase is an essential type II topoisomerase that introduces negative supercoils into DNA through an ATP-dependent mechanism (Gellert *et al*., 1976a; Higgins *et al*., 1978; Nöllmann *et al*., 2007); it can also relax negatively supercoiled DNA via an ATP-independent mechanism (Gellert *et al*., 1977; Higgins *et al*., 1978; Williams and Maxwell, 1999). The enzyme is composed of two copies of two subunits, GyrA and GyrB, giving it an A_2_B_2_ structure (Bates and Maxwell, 2005). Topoisomerase activity is required to eliminate the over-wound (positively supercoiled) and under-wound (negatively supercoiled) zones of the DNA template that are generated by transcription and DNA replication (Liu and Wang, 1987; Stracy et al., 2019). DNA gyrase relaxes the positively supercoiled DNA by introducing negative supercoils in an ATP-dependent manner (Ashley et al., 2017). Transcription and the associated disturbance to local DNA topology contribute to the architecture of the bacterial nucleoid by influencing the probability of DNA-DNA contacts between parts of the genome that border long transcription units that are heavily transcribed (Le and Laub, 2016). The changes to local DNA supercoiling that are caused by transcription and DNA replication also affect the activities of some transcription promoters (Ahmed et al., 2016; 2017; Chong et al., 2014; Dorman, 2019; Higgins, 2014; Rahmouni and Wells 1992; Rani and Nagaraja, 2019; Wu et al., 1988; Tobe et al., 1995). A large subset of promoters is sensitive to alterations in DNA supercoiling and to ensure appropriate gene expression, topoisomerases play an important role in maintaining supercoiling set points within a range that is tolerable by the cell (Cheung et al., 2003; Dorman and Dorman, 2016; Peter et al., 2004; Sutormin et al., 2019). The promoters of *gyrA* and *gyrB*, the genes that encode the A and B subunits, respectively, of gyrase, are stimulated by DNA relaxation (Menzel and Gellert, 1983; 1987; Straney et al., 1994; Unnirahman and Nagaraja, 1999). This is part of a mechanism that maintains DNA supercoiling homeostasis, keeping average DNA supercoiling values within the tolerable range (DiNardo et al., 1982; Dorman et al, 1989; Pruss et al., 1982; Raji et al., 1985; Richardson et al., 1988; Steck et al., 1984). As a corollary to this, the transcription of *topA*, the gene encoding DNA topoisomerase I (Topo I), is stimulated by negative supercoiling (Ahmed et al., 2016; Tse-Dinh and Beran, 1988). Topo I relaxes negatively supercoiled DNA using an ATP-independent type I mechanism (Dekker *et al*., 2002).

Several studies have shown that gene position on the chromosome is physiologically significant in bacteria (Bogue et al., 2020; Bryant et al., 2014; Gerganova et al., 2015; Scholz et al., 2019). In *Salmonella enterica* serovar Typhimurium, the *gyrA* and *gyrB* genes are widely separated on the genetic map of the circular chromosome: the *gyrB* gene is located close to the origin of chromosome replication, *oriC*, while *gyrA* is located near to the terminus region, Ter (McClelland et al., 2001). This arrangement closely resembles that seen in the model organism, *Escherichia coli* (Berlyn, 1998; Blattner et al., 1997). It has been proposed that the order of genes along each replichore in the bidirectionally replicated circular chromosome of *E. coli* correlates with the peak levels of expression of genes as a culture passes through each of the major stages of its growth cycle in batch culture (Sobetzko et al., 2012). DNA supercoiling plays an important role in the initiation of chromosome replication, so locating *gyrB* close to *oriC* is consistent with the gene location hypothesis. Bacteria emerging from lag phase and entering a period of rapid growth in exponential phase, experience a build up of negative DNA supercoiling and this stimulates the transcription of genes whose products support rapid growth (Colgan et al., 2018; Conter et al., 1997). Rapidly growing bacteria undergo multiple rounds of initiation of chromosome replication, so genes close to *oriC* are present in more copies per cell than those close to the terminus (Cooper and Helmstetter, 1968). GyrA and GyrB are required in equal amounts to form active DNA gyrase molecules, so the physical separation of *gyrA* from *gyrB* on the chromosome, and their organisation as independent transcription units, seem counterintuitive.

The genetically separated pattern of *gyrA* and *gyrB* gene location seen in *S*. Typhimurium and *E. coli* is not found universally among bacteria: many possess a *gyrBA* operon, although none appears to have a *gyrAB* operon. Examples of bacteria with a *gyrBA* setup include, *inter alia, Listeria monocytogenes* (Glaser et al., 2001), *Mycobacterium tuberculosis* (Cole et al., 1998) and *Staphylococcus aureus* (Baba et al., 2008). The operon arrangement appears to offer a number of advantages over the individual transcription unit model. Co-expression allows *gyrA* and *gyrB* to share the same promoter and the same transcription regulatory signals. Production of GyrA and GyrB from a common, bicistronic mRNA is likely to facilitate the establishment of equal amounts of each protein. The co-production of GyrA and GyrB might also be expected to enhance the efficiency of gyrase enzyme assembly. It should be noted that the *gyrA* and *gyrB* genes are only seen to be widely separated from one another on the unfolded, circular genetic map of *S*. Typhimurium: the genes may be brought into closer proximity in the folded chromosome within the nucleoid. Furthermore, in the tiny universe of the bacterium’s interior, the problem of gyrase assembly from GyrA and GyrB subunits produced from spatially separated mRNA molecules may be an insignificant one (Moffitt et al., 2016). We investigated this issue by building a derivative of *S*. Typhimurium with a *gyrBA* operon and comparing its physiology with that of the wild type.

## Results

### *Constructing a derivative of* S. *Typhimurium with a synthetic* gyrBA *operon*

A kanamycin resistance cassette, *kan*, was inserted adjacent to the *gyrA* gene in *S*. Typhimurium strain SL1344 to serve as a selectable marker (Experimental procedures). This *gyrA-kan* combination was amplified by PCR, leaving behind the transcription control signals of *gyrA*, and inserted immediately downstream of the *gyrB* gene, creating a *gyrBA* operon with an adjacent *kan* gene that was bordered by directly-repeated copies of the FRT sequence; the *kan* gene was then deleted by FLP-mediated site-specific recombination at the *frt* sites. A *kan* gene cassette, flanked by directly repeated *frt* sites, was used to replace the *gyrA* gene at the native *gyrA* location in the *gyrBA*-operon-containing strain; this *kan* cassette was then excised by FLP-mediated recombination. This process produced a derivative of SL1344 that had a *gyrBA* operon at the chromosomal position that is normally occupied by just *gyrB* and had no *gyrA* gene at the chromosomal site where this gene normally resides (Fig. 1). The whole genome sequence of this new strain was determined to ensure that no genetic changes, other than the desired ones, were present; none were detected.

**Fig. 1.**
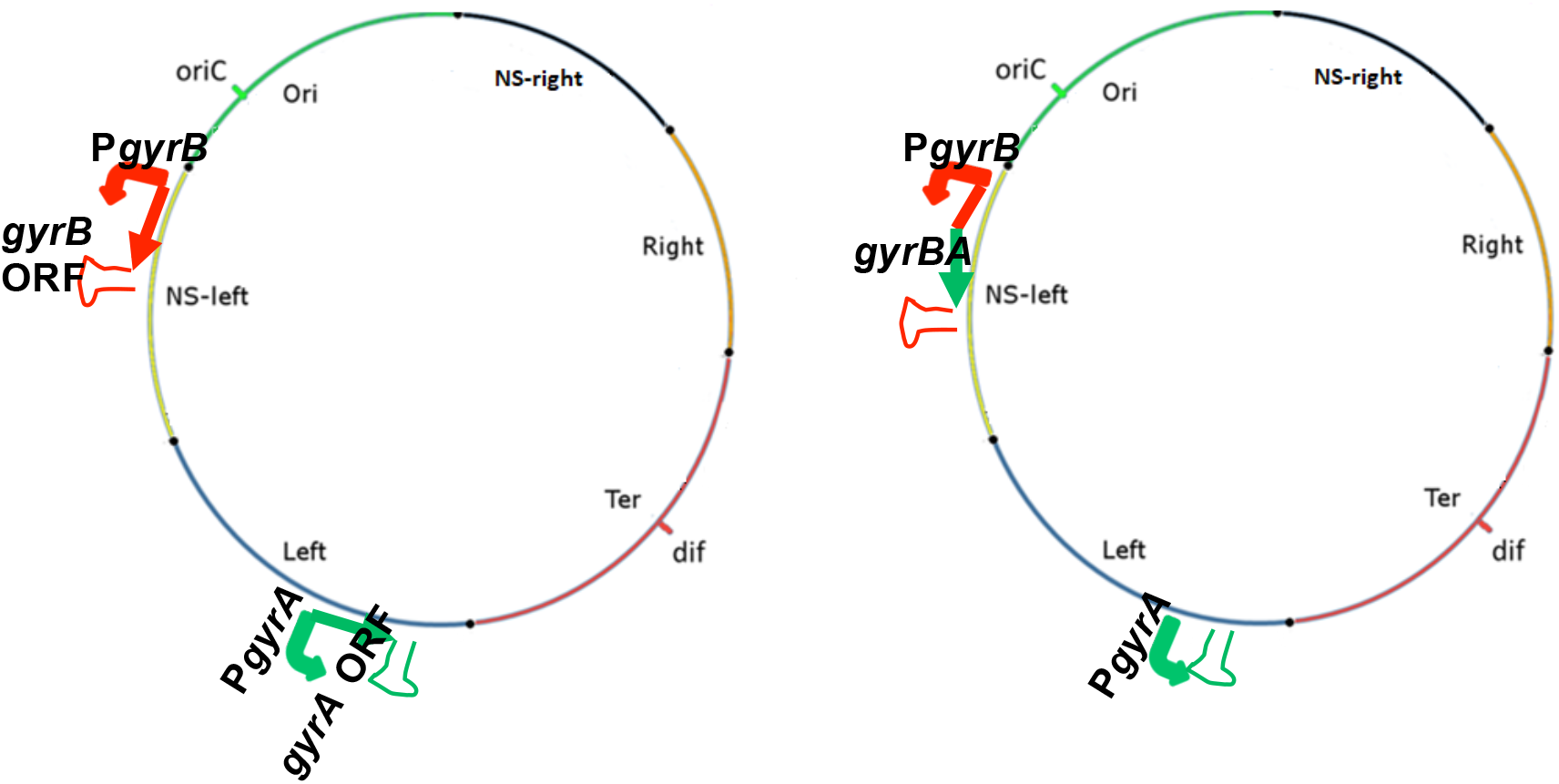
Construction of a derivative of *S*. Typhimurium strain SL1344 with a *gyrBA* operon. Chromosomal maps of the WT SL1344 and SL1344 *gyrBA* strains. Positions of *oriC, dif* and macrodomains are shown. Promoter, protein coding region (open reading frame, ORF) and the terminator of the genes of interest are shown and colour coded. The *gyrA* ORF is green and the *gyrB* promoter and ORF are red. Not to scale.

An attempt was also made to build a derivative of SL1344 with a *gyrAB* operon at the native location of the *gyrA* gene using a similar strategy to that used in constructing the synthetic *gyrBA* operon. Although the desired *gyrAB* operon was produced, it proved to be impossible to delete the second copy of *gyrB* from its native location. This suggested that the presence of the *gyrB* gene at its native *oriC*-proximal location is essential, even when a second copy of *gyrB* is present at another chromosomal position.

### *The growth characteristics of the* gyrBA *operon strain*

The growth kinetics of the wild type and the strain with the *gyrBA* operon were compared in batch liquid culture. Lysogeny broth (LB) cultures had identical growth curves when measured by plating and colony counting or by optical density measurements (Fig. S1A, S1B). Growth was also assessed in a minimal medium in an experiment that included low magnesium stress, an important environmental challenge that *S*. Typhimurium encounters in the macrophage vacuole during infection (Colgan et al., 2018). The wild type and the *gyrBA* operon strains were grown in minimal medium N with either 10 μM (low magnesium) or 10 mM (high magnesium) MgCl_2_. Once again, the two strains had identical growth kinetics (Fig. S1C).

### *Morphology of the strain with the* gyrBA *operon*

The identical growth characteristics of the strain with the *gyrBA* operon and the wild type, both in LB and in minimal medium, showed that producing DNA gyrase from an operon made no difference to the growth cycle and suggested that the cell cycle was unlikely to be altered either. Interference with the timing of major events in the cell cycle (initiation, replication fork passage and termination) can lead to delays in cell division, resulting in filamentation of the bacterial cell (Martin et al., 2020; Sharma and Hill, 1995). When we compared the morphologies of mid-exponential-phase cultures of the wild type and the *gyrBA* operon strain by light microscopy, no differences in the shapes of the cells or the frequency of cell filamentation were detected (Fig. S2). Taken together with the growth kinetic data, these findings showed that the operonic arrangement of *gyrA* and *gyrB* was well tolerated by *S*. Typhimurium.

### Sensitivity to gyrase-inhibiting antibiotics

The minimum inhibitory concentrations of gyrase-inhibiting antibiotics were compared for wild type SL1344 and SL1344 *gyrBA* (Fig. 2). Four drugs were tested: coumermycin and novobiocin are coumarins that target the GyrB subunit of DNA gyrase (Lewis et al., 1996) while nalidixic acid and ciprofloxacin are quinolones that target GyrA (Drlica and Zhao, 1997). The two strains were equally sensitive to the quinolones, but the SL1344 *gyrBA* strain was more resistant than SL1344 to novobiocin while SL1344 was more resistant than SL1344 *gyrBA* to coumermycin (Fig. 2). The reasons for the differential sensitivity patterns of the strains to the two classes of antibiotics, and for the opposing patterns of resistance to the two coumarins were not determined. However, the results indicated that producing the subunits of gyrase from a *gyrBA* operon resulted in coumarin MIC data that were not equivalent to those measured for the strain producing the subunits from physically separate genes.

**Fig. 2.**
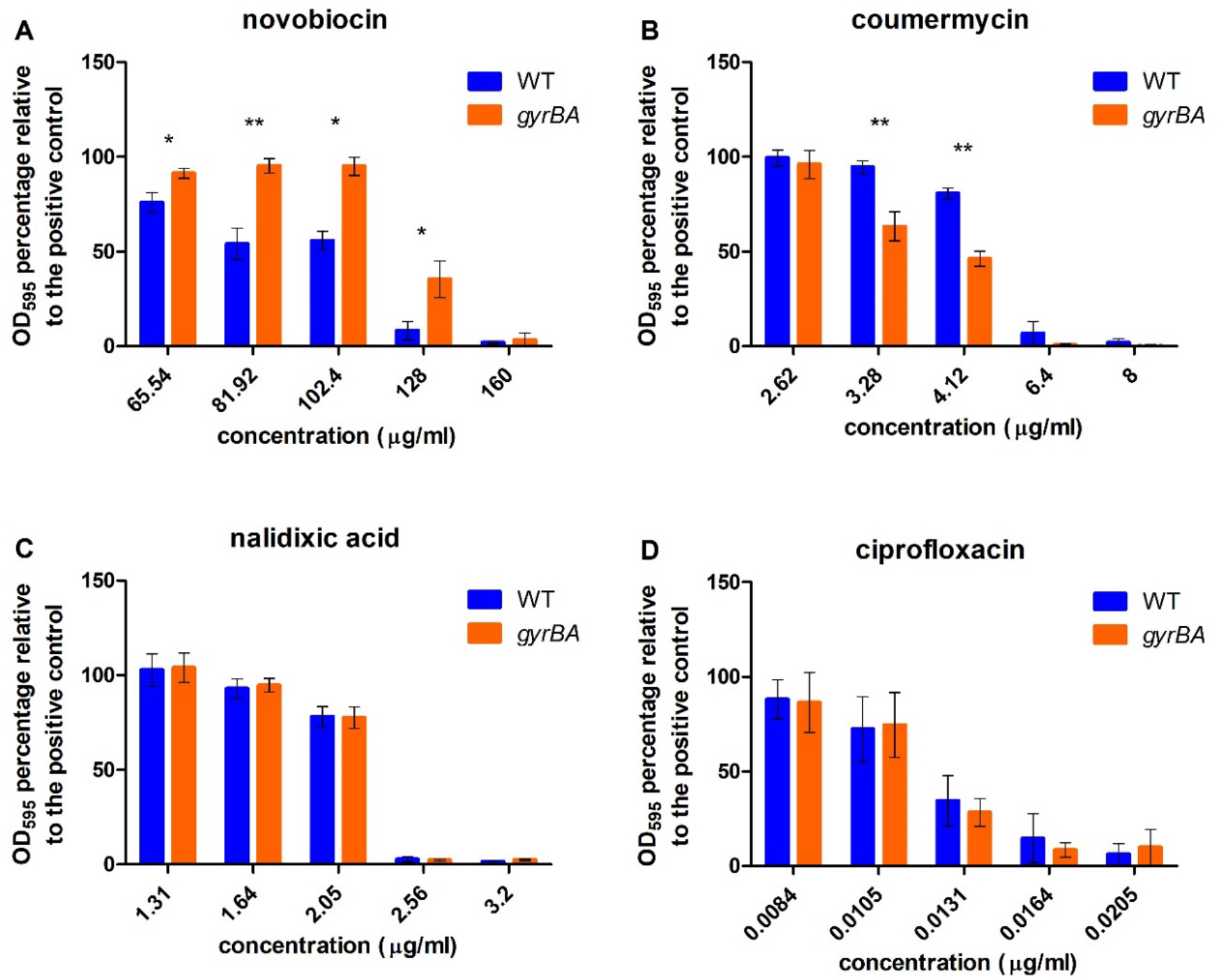
Minimum inhibitory concentrations of DNA gyrase-inhibiting antibiotics in the wild type SL1344 and SL1344 *gyrBA* strains. Cells were grown in a 96-well plate with 1:1.25 serially diluted antibiotics in LB broth for 18 h at 37°C and aeration. Cell density was measured by OD_600_. A. Percentage survival of the WT and SL1344 *gyrBA* in 65.54-160 µg/ml novobiocin. MIC_90_ of the WT = 128 µg/ml, MIC_90_ of SL1344 *gyrBA* = 160 µg/ml. B. Percentage survival of the WT and the *gyrBA* in 2.62-8 µg/ml coumermycin, MIC_90_ = 6.4 µg/ml. C. Percentage survival of the WT and the *gyrBA* in 1.31 – 3.2 µg/ml nalidixic acid, MIC_90_ = 2.56 µg/ml. D. Percentage survival of the WT and the *gyrBA* in 0.0084 – 0.0205 µg/ml ciprofloxacin, MIC_90_ = 0.0164 µg/ml. Error bars represent the standard deviation of at least three biological replicates. Significance was found by unpaired Student’s t-test, where * = P<0.05 and ** = P<0.01.

### Motility and competitive fitness measurements

The *gyrBA* operon strain was compared with the wild type to assess relative motility on agar plates and competitive fitness in liquid co-culture. The operonic strain showed a small, but statistically significant, reduction in motility compared to the wild type (Fig. 3A). The reasons for this were not determined and may reflect changes at one or more levels in the production and operation of the complex motility machinery of the bacterium. In contrast, the two strains were equally competitive when growing in LB (Fig. 3B). To perform the competition, the two strains were each marked genetically by insertion on a chloramphenicol resistance (*cat*) cassette that is located in the pseudogene *SL1483*. This *cat* insertion has a neutral effect on fitness and serves simply to allow the competing bacteria to be distinguished by selection on chloramphenicol-containing agar (Lacharme-Lora et al., 2019). The competitions were performed between a *cat*-marked wild type and the unmarked *gyrBA* operon strain and separately between a *cat*-marked *gyrBA* operon strain and the unmarked wild type (Fig. 3B). No differences in the competitive indices of the two strains were detected in either version of the competition.

**Fig. 3.**
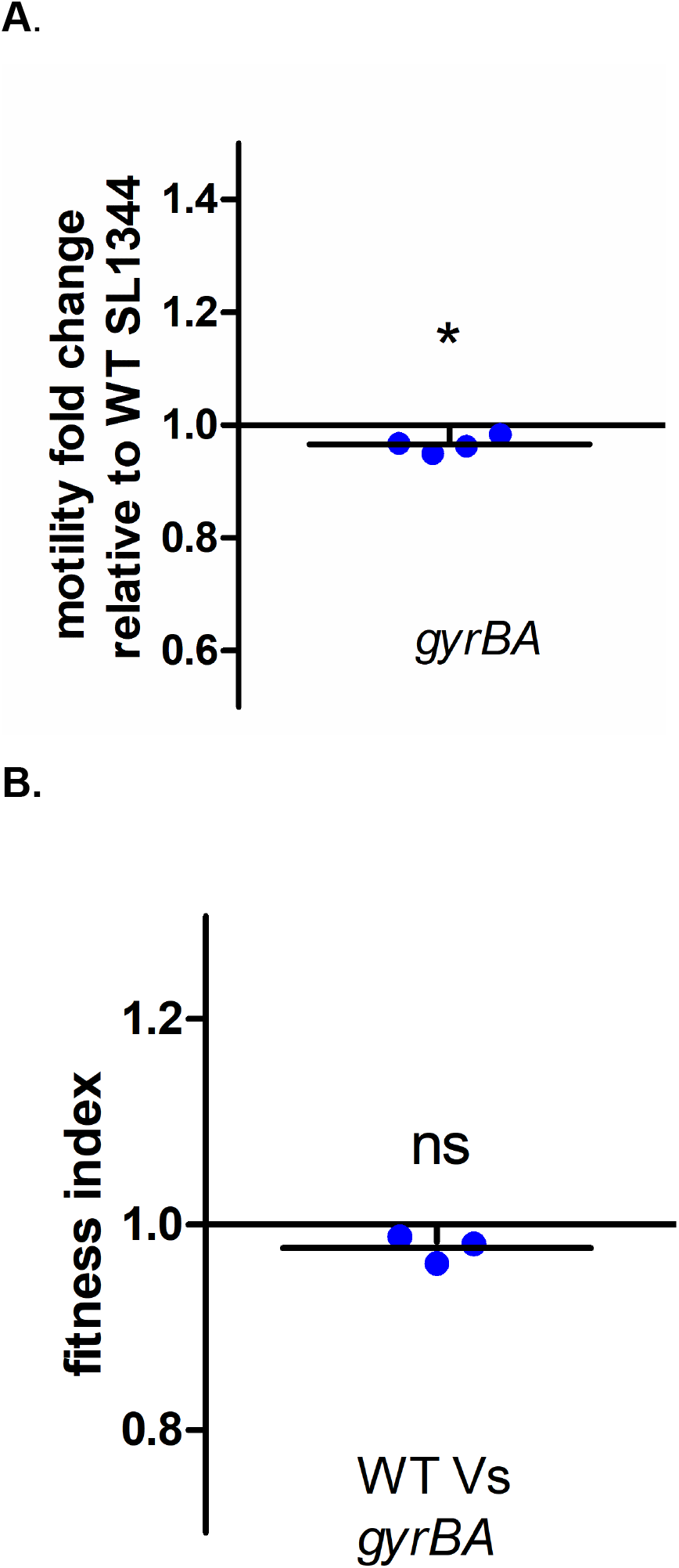
Motility and competitive fitness of strain SL1344 *gyrBA*. A. Diameters of swimming motility were measured after 5 h incubation at 37°C on soft 0.3% LB agar. The *gyrBA* strain is slightly, but statistically significantly, less motile than the WT. Values below 1 indicate that the strain is less motile than the WT. B) Fitness of the *gyrBA* strain was compared to the WT SL1344 in LB broth grown for 24 h with aeration at 37°C. Fitness index (f.i.) = 1 means that the competed strains were equally fit, f.i. < 1 indicates that the competitor strain is less fit than the WT, f.i. > 1 indicates that the competitor is more fit than the WT. The *gyrBA* and the WT were equally fit. Significance was determined by one sample T-test, where P<0.05.

### *Transcription of the separate and the operonic* gyr *genes*

The output of mRNA from the *gyrA* and *gyrB* genes was measured by quantitative PCR in wild type SL1344 and in SL1344 *gyrBA*, using the transcript of the *hemX* gene as a reference. (Expression of the *hemX* gene does not change under the growth conditions used here [Kröger *et al*., 2013]). Gyrase gene transcription in both strains was found to vary with growth cycle stage, with mRNA outputs being highest in early exponential phase (2-h time point) and lowest in stationary phase (Fig. 4). In the wild type, the *gyrA* gene (located near Ter, the terminus of chromosome replication) was expressed to a significantly higher level than *gyrB* (located close to *oriC*) at 2 h. This was interpreted as a reflection of the need to compensate for the effect of increased *gyrB* gene dosage relative to that of *gyrA* in rapidly growing cells. As the culture approached stationary phase, the levels of *gyrA* and *gyrB* transcripts equalised, in line with the convergence of *oriC*-proximal and Ter-proximal gene dosages (Fig. 4). The formation of the *gyrBA* operon eliminated the difference in *gyrB* and *gyrA* mRNA levels because each became part of the same bicistronic transcript and has adopted the expression profile of *gyrB* (Fig. 4).

**Fig. 4.**
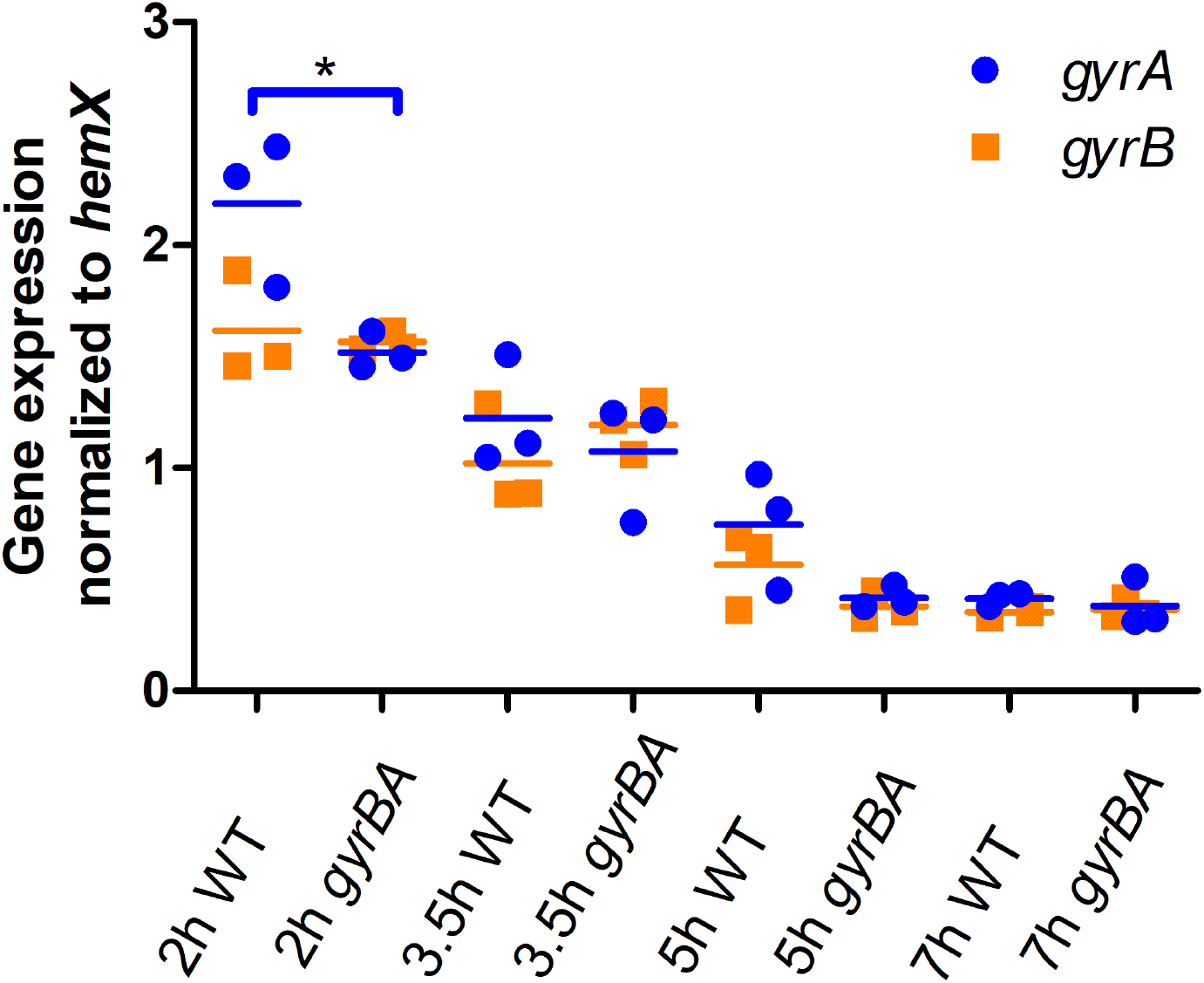
Expression of the *gyrA* and *gyrB* genes in wild type SL1344 (WT) and SL1344 *gyrBA* during growth in liquid culture. Cells were grown in LB broth at 37°C with aertion and samples were taken at 2 h, 3.5 h, 5 h and 7 h representing the lag, exponential, exponential-stationary transition and early stationary phases of growth, respectively. Transcription of *gyrA* and *gyrB* was measured and is reported relative to that of the *hemX* reference gene. Three biological replicates were used. Statistical significance was found by unpaired Student’s T-test, where P<0.05.

### *DNA supercoiling in the strain with a* gyrBA *operon*

The distributions of the topoisomers of the pUC18 reporter plasmid isolated from the wild type and the *gyrBA* operon strain were compared by electrophoresis in a chloroquine-agarose gel (Fig. 5). The cultures were grown in LB medium (Fig. 5A; S3A) or in minimal medium N with high or low concentrations of MgCl_2_ (Fig. 5B; S3B). In LB, the reporter plasmid was more relaxed in the *gyrBA* strain than in the wild type (Fig. 5A, S3B). Low-magnesium growth was used to mimic one of the stresses experienced by *Salmonella* in the macrophage vacuole. In the high MgCl_2_ control, the wild type and the *gyrBA* operon strain differed in their reporter plasmid distributions: DNA from the *gyrBA* strain was more negatively supercoiled than that from the wild type and showed a linking number difference (ΔLk) of - 3 (Fig. 5b, S3B). At the low MgCl_2_ concentration, the topoisomer distributions were more relaxed in both strains than in the high MgCl_2_ controls (ΔLk = +3). The reporter plasmid from the *gyrBA* operon strain was also more negatively supercoiled than that from the wild type, with the peak in its topoisomer distribution being approximately 2 linking numbers below that of the wild type (Fig. 5B, S3B). DNA relaxation occurs in *Salmonella* cells as they adjust to the macrophage vacuole and this forms part of the activation mechanism for the genes in the SPI-2 pathogenicity island (Cameron and Dorman, 2012; O Cróinín et al., 2006; Quinn et al., 2014). The products of these virulence genes protect the bacterium by inhibiting fusion of the vacuole with lysosomes (Garvis et al., 2001). We therefore monitored SPI-2 gene transcription in our two strains.

**Fig. 5.**
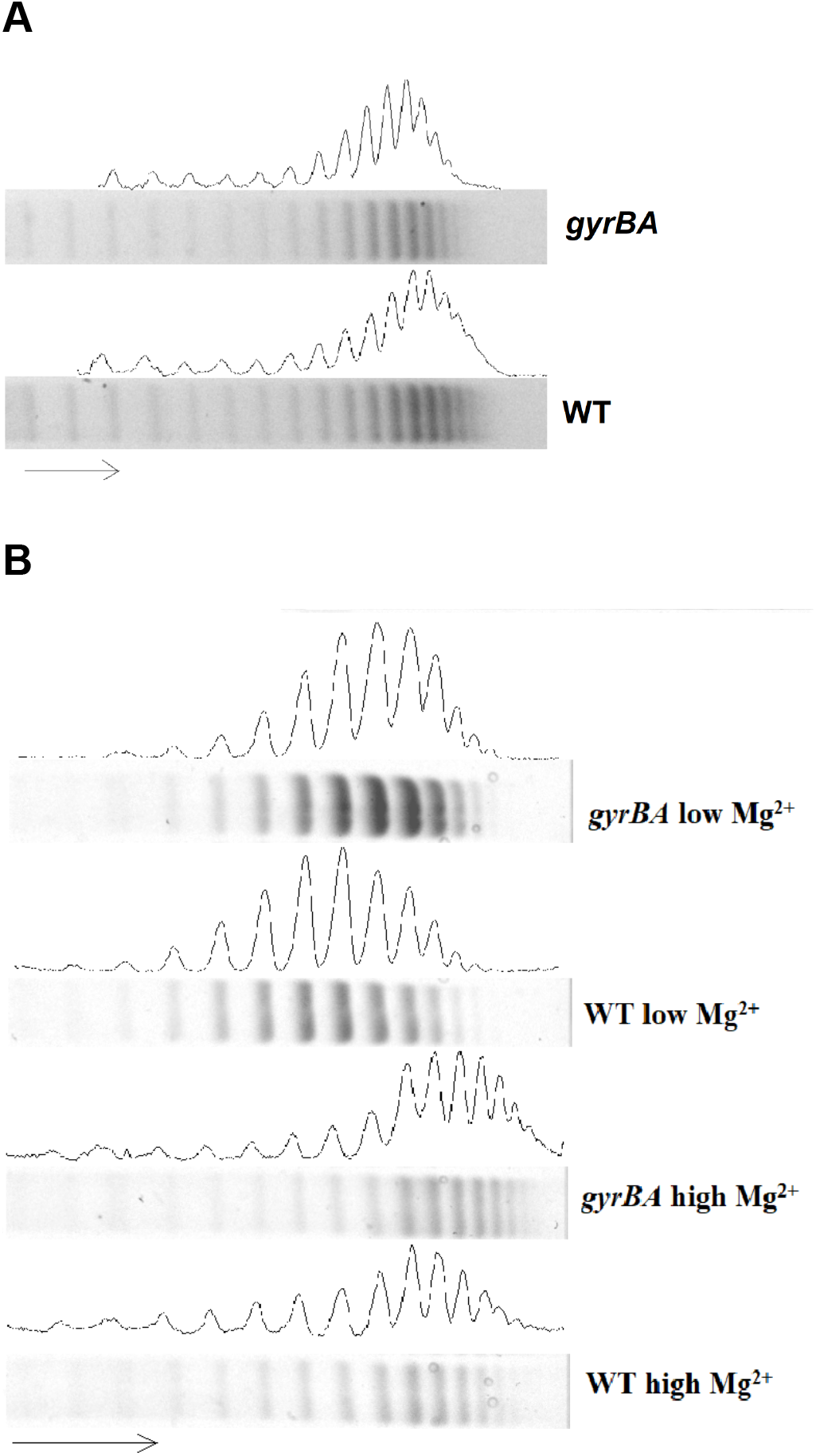
Reporter plasmid DNA supercoiling in SL1344 *gyrBA*. The pUC18 reporter plasmid was extracted from the WT and the SL1344 *gyrBA* strains at the stationary phase of growth and electrophoresed on a 0.8% agarose gel containing 2.5 µg/ml chloroquine.. The arrow shows the direction of migration, with the more supercoiled plasmid topoisomers at the right of the gel. A. Global DNA supercoiling pattern of the WT and the *gyrBA* strain when grown in LB. B. Global DNA supercoiling pattern of the WT and the *gyrBA* strain when grown in minimal medium N with high (10 mM) MgCl_2_ or low (10 µM) MgCl_2_. Sample lanes are supplemented with densitometry profiles that were generated with ImageJ. The analysis is representative of four biological replicates.

### *SPI-1 and SPI-2 gene expression in the* gyrBA *operon strain*

The SPI-1 and SPI-2 pathogenicity islands encode distinct type 3 secretion systems and effector proteins that are used to invade epithelial cells (SPI-1) or to survive in the intracellular vacuole (SPI-2) (Figueira and Holden, 2012; Hensel, 2000; van der Heijden and Finlay, 2012). Transcription of SPI-1 genes was monitored using a *gfp*^*+*^ reporter fusion under the control of the *prgH* promoter, P_*prgH*_, while a *gfp*^*+*^ fusion to the *ssaG* promoter (P_*ssaG*_) was used to monitor SPI-2 gene transcription. Wild type and *gyrBA* operon strains harbouring these fusions were grown in LB medium (Fig. 6A, 6B) or in minimal medium N, supplemented with high or low concentrations of MgCl_2_. (Fig. 6C-F). The cultures were grown with aeration at 37°C in 96-well plates and green fluorescence was measured throughout the growth cycle. The results obtained showed that in LB medium and in minimal medium N, SPI-1 transcription was indistinguishable between the wild type and the *gyrBA* operon strains (Fig. 6A, 6C, 6E). SPI-2 transcription was equivalent in both strains growing in LB (Fig. 6B) or in minimal medium with high MgCl_2_ (Fig. 6D). However, SPI-2 transcription occurred at reduced levels in the *gyrBA* strain in the later stages of the growth cycle under low magnesium conditions (Fig. 6F). These findings showed that when the subunits of gyrase are produced from an operon, rather than from separate *gyrA* and *gyrB* genes in their native chromosome locations, the normal expression profile of the SPI-2 virulence gene cluster is disrupted, but that this is conditional on growth in a low magnesium medium.

**Fig. 6.**
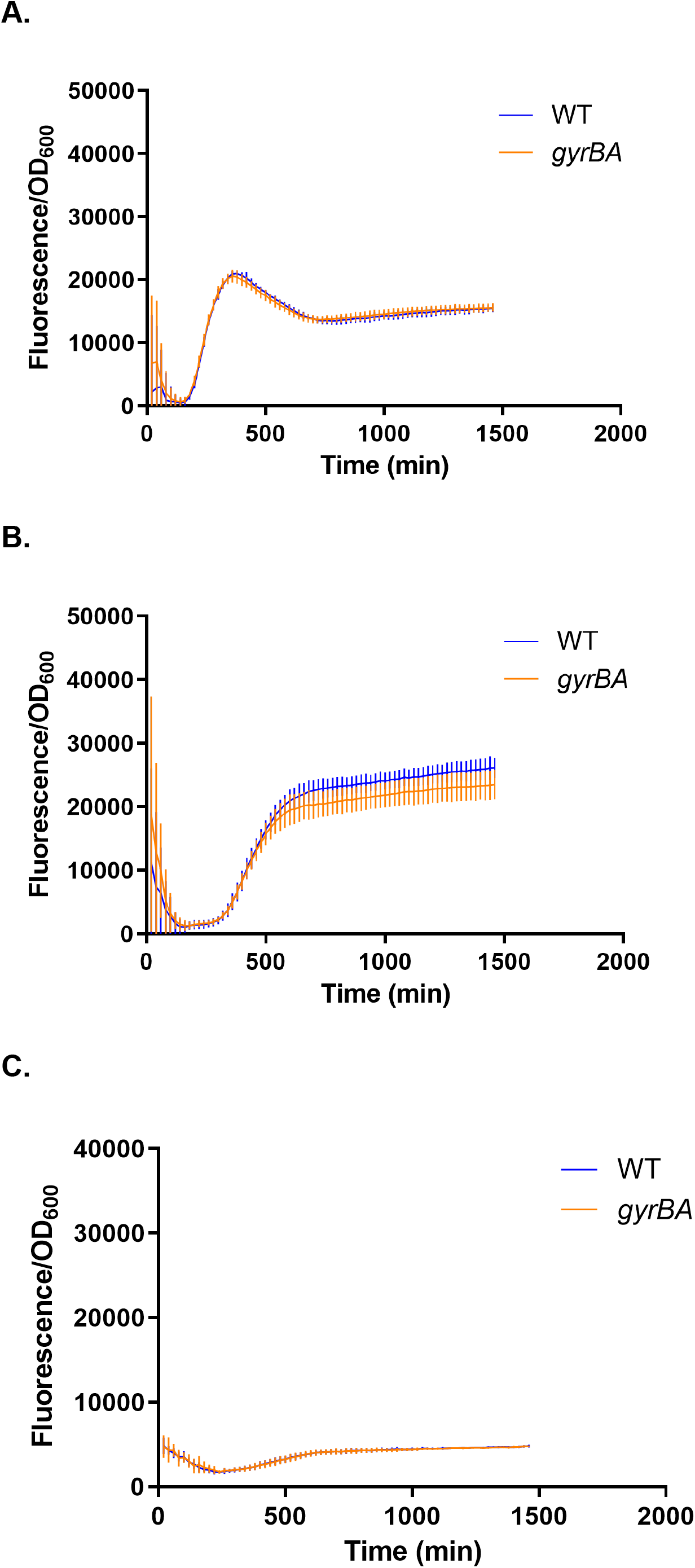
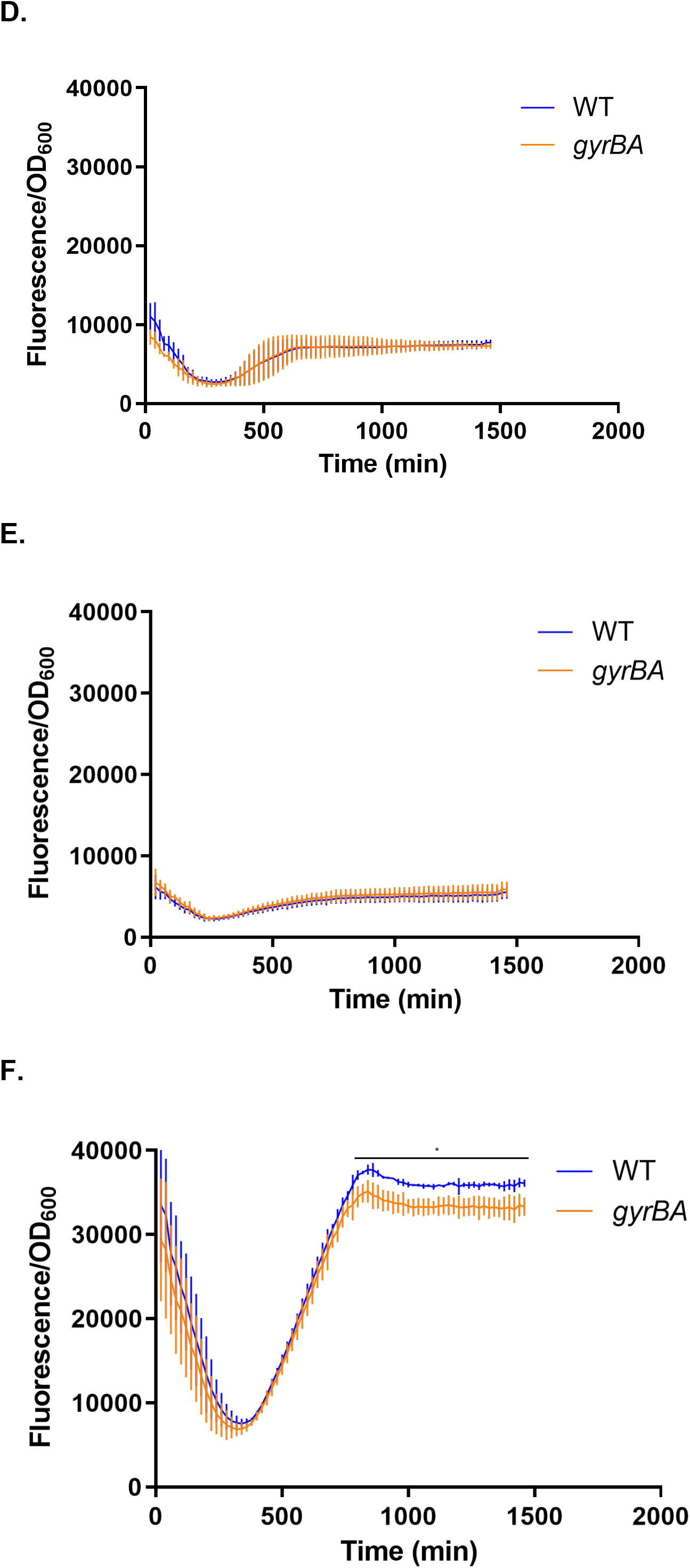
Expression of genes in the SPI-1 and SPI-2 pathogenicity islands in wild type SL1344 (WT) and SL1344 *gyrBA*. Expression of *gfp*^*+*^ reporter gene fusions was measured in the wild type and SL1344 *gyrBA* strains every 20 min over a 24-h. period. A. SPI-1 expression in the *gyrBA* strain was identical to that in the WT in LB. B. SPI-2 expression in the *gyrBA* strain was identical to that of the WT in LB. C. SPI-1 expression in minimal medium N with high MgCl_2_ concentration (10 mM) was repressed in both the WT and the *gyrBA* strain. D. SPI-2 expression in minimal medium N with high MgCl_2_ concentration was repressed in both the WT and the *gyrBA* strains. E. SPI-1 expression in minimal medium N with a low MgCl_2_ concentration (10 μM) was repressed in both the WT and the *gyrBA* strains. F. SPI-2 expression in minimal medium N with low MgCl_2_ concentration was lower in the *gyrBA* strain than in the WT at the stationary phase of growth. All plots are the results of at least three biological replicates; error bars represent the standard deviation. Statistical significance was found by Student’s unpaired T-test, where P<0.05.

### *Impact of the gyrase operon on cell infection by* Salmonella

The abilities of the wild type and the *gyrBA* operon strains to invade and to replicate in cultured mammalian cells were compared. Bacteria, grown to mid-exponential phase to promote SPI-1 gene expression, were used to infect RAW264.7 macrophage. When intracellular bacteria were then enumerated post-invasion, fewer of the *gyrBA* operon strain cells were detected than wild type cells at and after the 16-h time point (Fig.7). This reduction in bacterial survival correlated with the diminished SPI-2 expression detected in the *gyrBA* operon strain in low Mg^2+^, a macrophage relevant condition.

## Discussion

The genes in *Salmonella* that encode DNA gyrase, *gyrA* and *gyrB*, are located at the opposite ends of the left replichore of the chromosome (Fig. 1). In contrast, many other bacteria, such as *Listeria monocytogenes, Staphylococcus aureus* and *Mycobacterium tuberculosis*, possess a *gyrBA* operon (Baba et al., 2008; Cole et al., 1998; Glaser et al., 2001). As a first step in assessing the significance of the stand-alone gyrase gene arrangement versus the operon model, we constructed a derivative of *S*. Typhimurium with a *gyrBA* operon at the chromosomal location that is normally occupied by *gyrB* alone, while removing the individual *gyrA* gene from its native position in the genome. This strain, with the *gyrA* and *gyrB* genes transcribed from a common promoter (P_*gyrB*_) and located close to the origin of chromosomal replication, had normal growth characteristics and cell morphology. The deletion of the (now redundant) *gyrA* gene copy from its native position near the terminus of chromosome replication was tolerated well. In contrast, it proved to be impossible to delete the *gyrB* gene from its native location in a strain that had a *gyrAB* operon at the chromosomal position that is normally occupied by *gyrA* alone. These findings show that there are limits to the rearrangements of *gyrA* and *gyrB* genes that *Salmonella* will tolerate.

Although the production of DNA gyrase from an operon was well tolerated by *S*. Typhimurium, the operon strain differed from the wild type in a number of phenotypic characteristics. Subtle differences in sensitivities to coumarin antibiotics, but not quinolones, distinguished the operon strain from the wild type (Fig. 2). The reasons for the different responses to coumarins were not determined and may have involved indirect effects of operonic gyrase production on processes involved in drug uptake. A modest decrease in competitive fitness in the operon strain hinted at impacts on physiology. In light of the fact that *S*. Typhimurium is a facultative intracellular pathogen, it was interesting to note a difference in the expression of the horizontally acquired SPI-2 pathogenicity island. The *gyrBA* operon strain expressed SPI-2 less well (Fig. 6) and survived less well than the wild type in the macrophage (Fig. 7), whose *Salmonella*-containing vacuole is a stressful, low magnesium environment where SPI-2 plays a key protective role (Figueira and Holden, 2012; Hensel, 2000; van der Heijden and Finlay, 2012). The *gyrBA* operon strain maintained its DNA in a less relaxed state in a low-magnesium environment (Fig. 5). Since DNA relaxation contributes to full expression of SPI-2 genes (Cameron and Dorman, 2012; Quinn et al., 2014) and DNA in *S*. Typhimurium becomes relaxed when the bacterium is in the macrophage (O Cróinín et al., 2006), this may explain the poorer transcription of *ssaG* that was seen in low magnesium growth (Fig. 6).

**Fig. 7.**
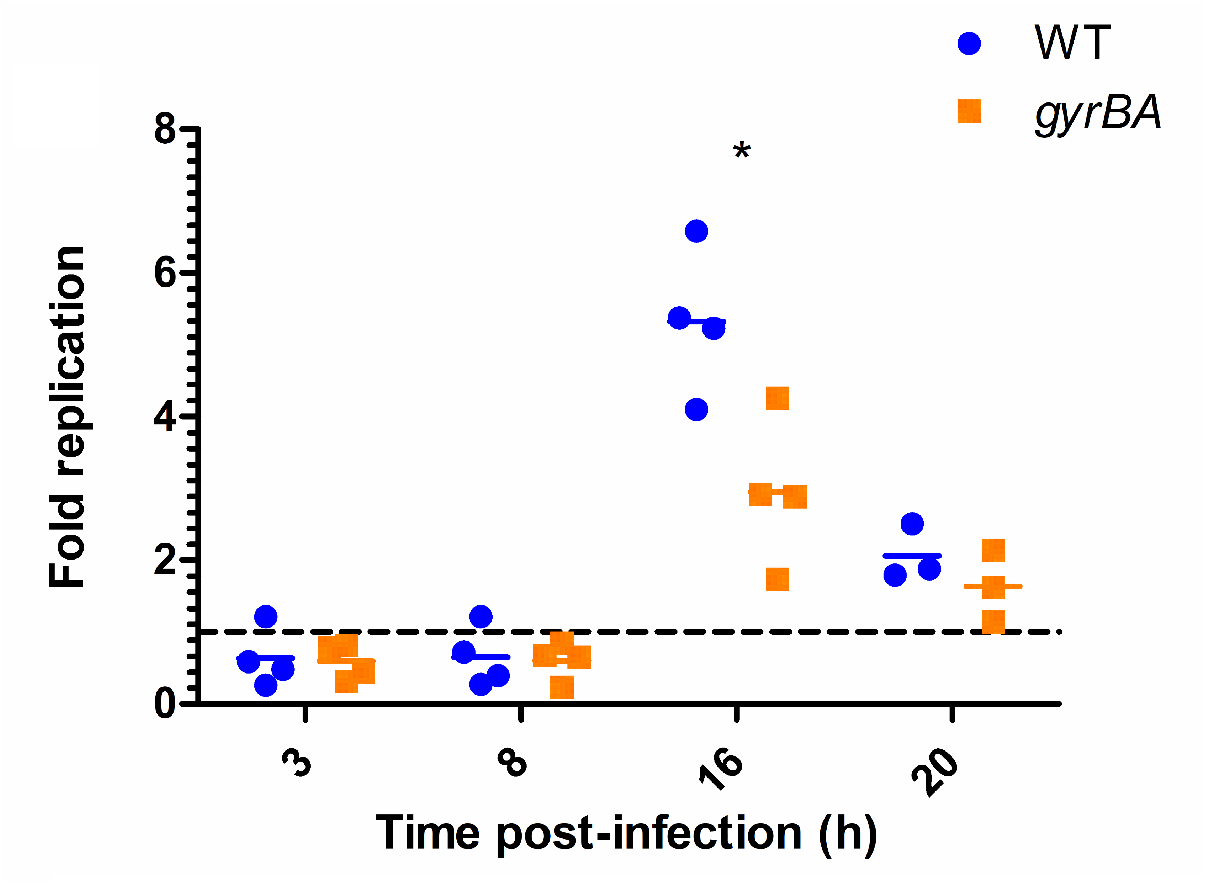
SPI-1-mediated entry and survival of the WT and SL1344 *gyrBA* strain in cultured RAW264.7 macrophage cells. Cells were infected with SPI-1-induced bacteria, grown to mod-exponential phase to promote SPI-1-mediated invasion. Survival and replication were measured by enumerating colony forming units (CFUs) at 3 h, 8 h, 16 h and 20 h post-infection. Fold replication represents the number of CFUs recovered at a particular time point divided by the CFU number ‘at 1 h. Mean and individual replicates are shown. Significance at 16 h was found by unpaired Student’s T-test, where P<0.05.

We have shown experimentally that there is no absolute barrier to the organisation of the *gyrA* and *gyrB* genes as a *gyrBA* operon in *Salmonella*. Furthermore, there are many examples of naturally occurring *gyrBA* operons among bacterial species. Why is this arrangement not found universally? Sharing a common promoter, common transcriptional regulatory features and a common chromosomal location would appear to offer the advantages of coordinated gene expression (Price et al., 2005) and physical co-production of protein products that will need to combine with a fixed stoichiometry to form an active product (Dandekar et al., 1998; Pal and Hurst, 2004; Swain, 2004). Indeed, the coupling of transcription and translation in prokaryotes may aid the production of operon-encoded proteins that are required in stoichiometric amounts (Li et al., 2014; Rocha, 2008). Often, but not invariably, operons are composed of genes that contribute to a common pathway (de Daruvar et al., 2002; Lawrence and Roth, 1996; Price et al., 2006; Rogozin et al., 2002) and that is true of the *gyrBA* operon. Colocation of genes within an operon facilitates their collective translocation via horizontal gene transfer, allowing them to replace lost or mutated copies in the recipient cell (Lawrence and Roth, 1996). According to this “selfish operon” hypothesis, this creates a selective pressure for the maintenance of an operon structure. However, since loss of either *gyrA* or *gyrB* is lethal, the gyrase operon may be one of the exceptions to the selfish operon rule, because gyrase-deficient recipients cannot exist.

We conducted a survey of gyrase gene locations in bacteria to assess the frequency of the stand-alone arrangement seen in *Salmonella* and the *gyrBA* operon arrangement seen in other species (Experimental procedures). We were unable to find any examples of bacteria that naturally possess a *gyrAB* operon. The results of the survey are shown in Table 1, where bacteria are grouped according to their gyrase gene arrangement, using *oriC* as a reference point. Fig. 8 shows a phylogenetic tree summarising the occurrence of different gyrase gene arrangements among bacteria from the four groups listed in Table 1.

**Table 1.**
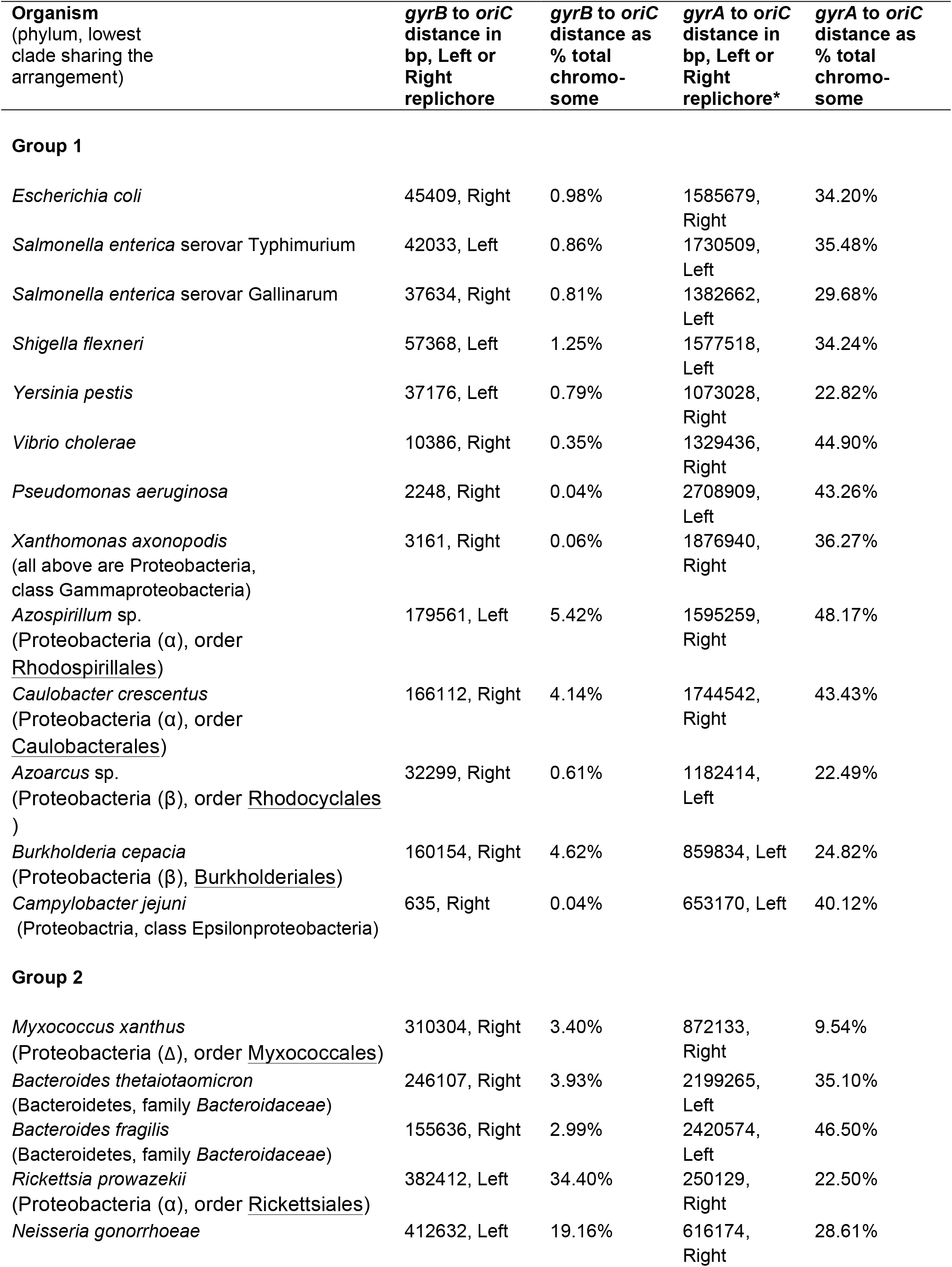
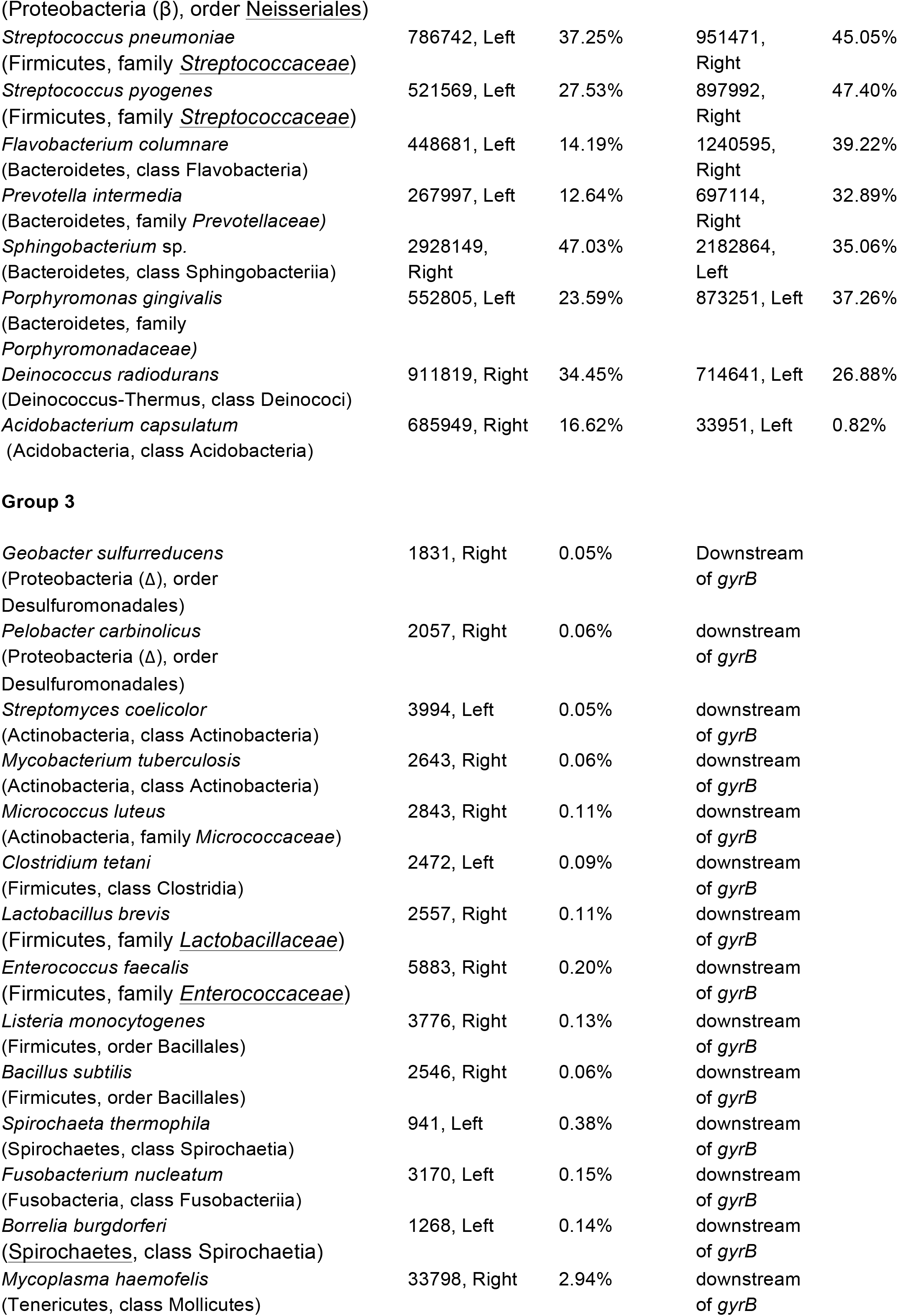
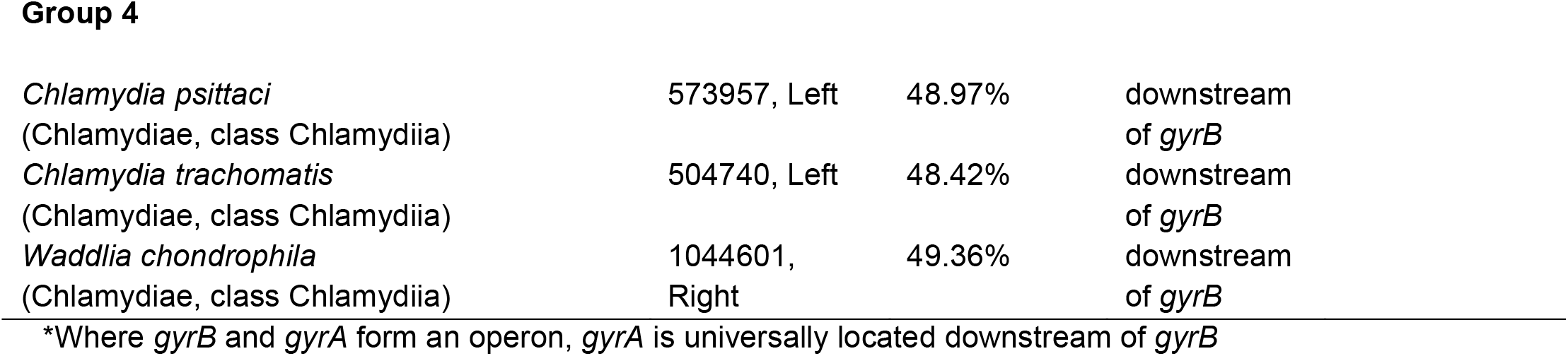
Relative positions of *gyrB* and *gyrA* across bacterial species

| Organism<br>(phylum, lowest<br>clade sharing the<br>arrangement) | *gyrB* to *oriC*<br>distance in<br>bp, Left or<br>Right<br>replichore | *gyrB* to *oriC*<br>distance as<br>% total<br>chromo-<br>some | *gyrA* to *oriC*<br>distance in<br>bp, Left or<br>Right<br>replichore* | *gyrA* to *oriC*<br>distance as<br>% total<br>chromo-<br>some |
| --- | --- | --- | --- | --- |
| **Group 1** |  |  |  |  |
| *Escherichia coli* | 45409, Right | 0.98% | 1585679,<br>Right | 34.20% |
| *Salmonella enterica* serovar Typhimurium | 42033, Left | 0.86% | 1730509,<br>Left | 35.48% |
| *Salmonella enterica* serovar Gallinarum | 37634, Right | 0.81% | 1382662,<br>Left | 29.68% |
| *Shigella flexneri* | 57368, Left | 1.25% | 1577518,<br>Left | 34.24% |
| *Yersinia pestis* | 37176, Left | 0.79% | 1073028,<br>Right | 22.82% |
| *Vibrio cholerae* | 10386, Right | 0.35% | 1329436,<br>Right | 44.90% |
| *Pseudomonas aeruginosa* | 2248, Right | 0.04% | 2708909,<br>Left | 43.26% |
| *Xanthomonas axonopodis*<br>(all above are Proteobacteria,<br>class Gammaproteobacteria) | 3161, Right | 0.06% | 1876940,<br>Right | 36.27% |
| *Azospirillum* sp.<br>(Proteobacteria (α), order<br><u>Rhodospirillales</u>) | 179561, Left | 5.42% | 1595259,<br>Right | 48.17% |
| *Caulobacter crescentus*<br>(Proteobacteria (α), order<br><u>Caulobacteriales</u>) | 166112, Right | 4.14% | 1744542,<br>Right | 43.43% |
| *Azoarcus* sp.<br>(Proteobacteria (β), order <u>Rhodocyclales</u><br>) | 32299, Right | 0.61% | 1182414,<br>Left | 22.49% |
| *Burkholderia cepacia*<br>(Proteobacteria (β), <u>Burkholderiales</u>) | 160154, Right | 4.62% | 859834, Left | 24.82% |
| *Campylobacter jejuni*<br>(Proteobacteria, class Epsilonproteobacteria) | 635, Right | 0.04% | 653170, Left | 40.12% |
| **Group 2** |  |  |  |  |
| *Myxococcus xanthus*<br>(Proteobacteria (Δ), order <u>Myxococcales</u>) | 310304, Right | 3.40% | 872133,<br>Right | 9.54% |
| *Bacteroides thetaiotaomicron*<br>(Bacteroidetes, family *Bacteroidaceae*) | 246107, Right | 3.93% | 2199265,<br>Left | 35.10% |
| *Bacteroides fragilis*<br>(Bacteroidetes, family *Bacteroidaceae*) | 155636, Right | 2.99% | 2420574,<br>Left | 46.50% |
| *Rickettsia prowazekii*<br>(Proteobacteria (α), order <u>Rickettsiales</u>) | 382412, Left | 34.40% | 250129,<br>Right | 22.50% |
| *Neisseria gonorrhoeae* | 412632, Left | 19.16% | 616174,<br>Right | 28.61% |
| (Proteobacteria (β), order Neisseriales) |  |  |  |  |
| *Streptococcus pneumoniae*<br>(Firmicutes, family <u>Streptococcaceae</u>) | 786742, Left | 37.25% | 951471, Right | 45.05% |
| *Streptococcus pyogenes*<br>(Firmicutes, family <u>Streptococcaceae</u>) | 521569, Left | 27.53% | 897992, Right | 47.40% |
| *Flavobacterium columnare*<br>(Bacteroidetes, class Flavobacteria) | 448681, Left | 14.19% | 1240595, Right | 39.22% |
| *Prevotella intermedia*<br>(Bacteroidetes, family <u>Prevotellaceae</u>) | 267997, Left | 12.64% | 697114, Right | 32.89% |
| *Sphingobacterium* sp.<br>(Bacteroidetes, class Sphingobacteriia) | 2928149, Right | 47.03% | 2182864, Left | 35.06% |
| *Porphyromonas gingivalis*<br>(Bacteroidetes, family <u>Porphyromonadaceae</u>) | 552805, Left | 23.59% | 873251, Left | 37.26% |
| *Deinococcus radiodurans*<br>(Deinococcus-Thermus, class Deinococi) | 911819, Right | 34.45% | 714641, Left | 26.88% |
| *Acidobacterium capsulatum*<br>(Acidobacteria, class Acidobacteria) | 685949, Right | 16.62% | 33951, Left | 0.82% |

**Group 3**
|  |  |  |  |
| --- | --- | --- | --- |
| *Geobacter sulfurreducens*<br>(Proteobacteria (Δ), order Desulfuromonadales) | 1831, Right | 0.05% | Downstream of *gyrB* |
| *Pelobacter carbinolicus*<br>(Proteobacteria (Δ), order Desulfuromonadales) | 2057, Right | 0.06% | downstream of *gyrB* |
| *Streptomyces coelicolor*<br>(Actinobacteria, class Actinobacteria) | 3994, Left | 0.05% | downstream of *gyrB* |
| *Mycobacterium tuberculosis*<br>(Actinobacteria, class Actinobacteria) | 2643, Right | 0.06% | downstream of *gyrB* |
| *Micrococcus luteus*<br>(Actinobacteria, family <u>Micrococcaceae</u>) | 2843, Right | 0.11% | downstream of *gyrB* |
| *Clostridium tetani*<br>(Firmicutes, class Clostridia) | 2472, Left | 0.09% | downstream of *gyrB* |
| *Lactobacillus brevis*<br>(Firmicutes, family <u>Lactobacillaceae</u>) | 2557, Right | 0.11% | downstream of *gyrB* |
| *Enterococcus faecalis*<br>(Firmicutes, family <u>Enterococcaceae</u>) | 5883, Right | 0.20% | downstream of *gyrB* |
| *Listeria monocytogenes*<br>(Firmicutes, order Bacillales) | 3776, Right | 0.13% | downstream of *gyrB* |
| *Bacillus subtilis*<br>(Firmicutes, order Bacillales) | 2546, Right | 0.06% | downstream of *gyrB* |
| *Spirochaeta thermophila*<br>(Spirochaetes, class Spirochaetia) | 941, Left | 0.38% | downstream of *gyrB* |
| *Fusobacterium nucleatum*<br>(Fusobacteria, class Fusobacteriia) | 3170, Left | 0.15% | downstream of *gyrB* |
| *Borrelia burgdorferi*<br>(Spirochaetes, class Spirochaetia) | 1268, Left | 0.14% | downstream of *gyrB* |
| *Mycoplasma haemofelis*<br>(Tenericutes, class Mollicutes) | 33798, Right | 2.94% | downstream of *gyrB* |

Group 4
|  |  |  |  |
| --- | --- | --- | --- |
| *Chlamydia psittaci*<br>(Chlamydiae, class Chlamydiia) | 573957, Left | 48.97% | downstream<br>of *gyrB* |
| *Chlamydia trachomatis*<br>(Chlamydiae, class Chlamydiia) | 504740, Left | 48.42% | downstream<br>of *gyrB* |
| *Waddlia chondrophila*<br>(Chlamydiae, class Chlamydiia) | 1044601,<br>Right | 49.36% | downstream<br>of *gyrB* |

\*Where *gyrB* and *gyrA* form an operon, *gyrA* is universally located downstream of *gyrB*

**Fig. 8.**
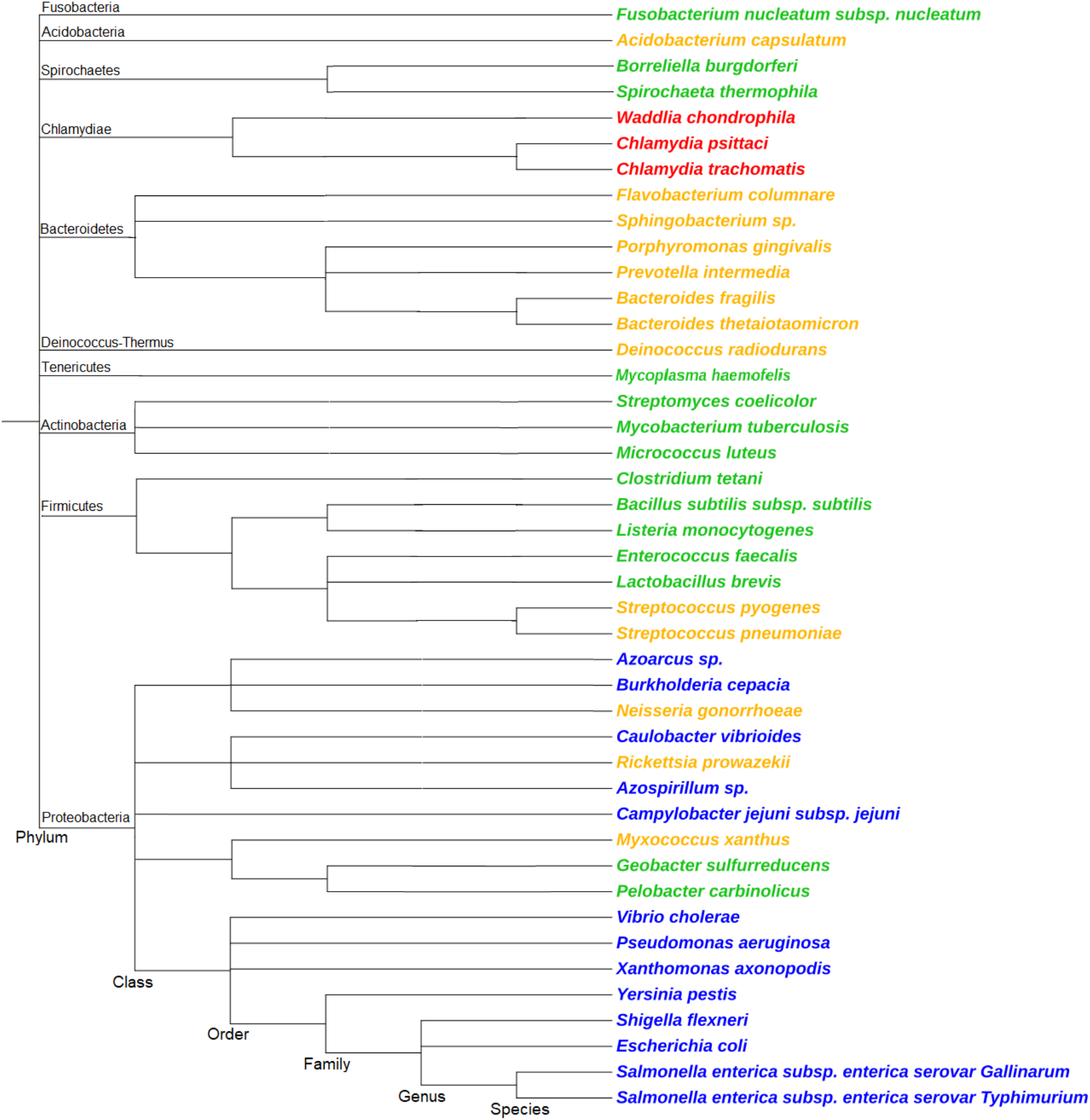
Phylogenetic tree of bacteria that belong to different groups based on their *gyrA* and *gyrB* arrangement. The phylogenetic tree was built in phyloT, a phylogenetic tree generator based on NCBI taxonomy (Letunic & Bork, 2019). Each of the four groups (see Table 1) of *gyrA* and *gyrB* arrangements is indicated by colour. Group 1, blue: *gyrA* and *gyrB* are at separate locations, with a conserved genetic environment 5’ to *gyrB*. Group 2, orange: *gyrA* and *gyrB* are at separate locations, with a non-conserved genetic environment 5’ to *gyrB*. Group 3, green: *gyrBA* operon, conserved genetic environment 5’ to *gyrB*. Group 4, red: *gyrBA* operon, non-conserved genetic environment 5’ to *gyrB*. Phyla names are indicated.

Inversions of DNA between the left and the right replichores were seen frequently and these followed no obvious patterns. This is in agreement with the previous finding that, while distance to the origin is highly conserved, inversions of genes around the Ter region of a chromosome are frequent and well tolerated between *E. coli* and *Salmonella* (Alokam *et al*., 2002). Various relative arrangements of *gyrA* and *gyrB* were observed and subdivided into four groups: Group 1 had *gyrB* and *gyrA* positioned separately, with *gyrB* near *oriC*; Group 2 had *gyrB* and *gyrA* positioned separately, with the position of *gyrB* being variable; Group 3 had *gyrB* and *gyrA* arranged as a *gyrBA* operon in the immediate vicinity of *oriC*; Group 4 had *gyrB* and *gyrA* arranged as a *gyrBA* operon at a distance from *oriC*. The arrangements of *gyrA* and *gyrB* genes were categorised into the four Groups not only according to the relative positions of *gyrA* and *gyrB* but also according to the degree of conservation of the genetic environment of *gyrB*.

Table 1 suggests that all members of the class gamma-proteobacteria (phylum Proteobacteria), including *E. coli* and *Salmonella*, some alpha-proteobacteria, beta-proteobacteria and epsilon-proteobacteria are in Group 1. Group 2 contains some alpha-, beta- and delta-proteobacteria, members of the family *Streptococcaceae* (order Lactobacillales, phylum Firmicutes); members of the class Flavobacteriia; multiple members of the phylum Bacteroidetes; Acidobacteria and *Deinococcus radiodurans*. Group 3 was composed of members of the phylum Actinobacteria, classes Clostridia and Bacilli (phylum Firmicutes), family *Enterococcaceae* and family *Lactobacillaceae* (order Lactobacillales, phylum Firmicutes) order Fusobacteria (phylum Fusobacteria) and phylum Terenicutes. Finally, Group 4 consisted of members of the phylum Chlamydiae. There is perhaps more variation within Group 4, but this was not detected using the method employed here. *Mycoplasma* is an anomaly of Group 3, since not all its species clearly belong to this group. Some *Mycoplasma* possess the expected conserved genes 5’ to *gyrB*, but not in its immediate vicinity. However, the orientation of genes 5’ to *gyrB* remains favourable for the initiation of its transcription, therefore, *Mycoplasma* is placed in Group 3. It was clear from the analysis that members of the same taxonomic rank do not necessarily have to belong to the same Group, especially in diverse phyla. For example, both Group 2 and Group 3 arrangements are present within the Firmicutes. Moreover, both arrangements are present within the order Lactobacillales alone. Some less diverse phyla such as Fusobacteria and Chlamydiae belong to only one Group. No variation was found within families.

It was difficult to conclude if given taxons were enriched in particular groups in Table 1, so a phylogenetic tree was plotted that included all of the bacteria in the table (Fig. 8). The tree was constructed using the phylogenetic tree generator phyloT, based on NCBI taxonomy (Letunic & Bork, 2019) and the positions of the branches were manually reviewed with the aid of the NCBI taxonomy browser. It is apparent that one Group can be present in multiple unrelated phyla and that one phylum can contain members of several Groups, illustrating a high level of diversity of *gyrA* and *gyrB* chromosomal arrangements. However, certain patterns are discernable. Bacteria from Group 1 are exclusively located in phylum Proteobacteria. All the members of phylum Bacteroidetes that were investigated belong to Group 2, but other phyla can also contain some members of Group 2. The Group 3 arrangement occurs with high frequency in the superphylum Terrabacteria (Firmicutes, Tenericutes, Actinobacteria, Deinococcus), although this arrangement can be encountered elsewhere too. Finally, all of the tree members of Group 4 shown belong to the phylum Chlamydiae. No other phylum was found to contain bacteria of Group 4, but the existence of the Group 4 arrangement outside of the Chlamydiae cannot be ruled out. It is also difficult to draw clear parallels between the lifestyle of an organism and the Group to which it belongs, since bacteria of various lifestyles can be members of the same Group. The analysis presented here is indicative rather than exhaustive: it is possible that further sampling will broaden the existing Groups and reveal further details.

We found no examples of a naturally occurring *gyrAB* operon in bacteria. Nevertheless, we attempted to construct one in *S*. Typhimurium. Although *gyrB* could be inserted successfully downstream of *gyrA*, attempts to delete the *gyrB* copy from its native locus failed repeatedly. The strong impediment to deleting *gyrB* from its original locus suggests the particular importance of the position of this gene. What are the constraints on removing *gyrB* from its natural location on the S. Typhimurium chromosome and why is *gyrA* free from such constraints? The *gyrB* and *gyrA* genes respond differently to treatment with DNA gyrase inhibitors (Neumann & Quiñones, 1997). In particular, the expression of both *gyrA* and *gyrB* can be induced by coumarins (inhibitors of the GyrA subunit), while only *gyrA* is induced by quinolones (inhibitors of the GyrB subunit). This is despite DNA relaxation occurring in response to both classes of gyrase-inhibiting drugs. This suggests that *gyrB* differs from *gyrA* in its sensitivity to DNA relaxation. Another characteristic that is different between these genes is the degree of sequence conservation in the regulatory regions. FIS regulation is important for *gyrA* and *gyrB* of in both *E. coli* and *Salmonella*. FIS regulatory binding sites are located upstream from the ORFs of both genes (Schneider *et al*., 1999; Keane & Dorman, 2003). However, the 5’ region of *gyrA* is significantly diverged between these bacteria, while the 5’ region of *gyrB* is, in contrast, highly conserved (Keane & Dorman, 2003). Various mutual arrangements of *gyrA* and *gyrB* were found among bacteria (Table 1), but the relative position of *gyrB* seems to be more conserved than that of *gyrA*.

When the immediate genetic environment of both genes in bacteria listed in Table 1 was studied, one distinct pattern was found – homologues of *dnaA* (encoding chromosomal replication initiation protein DnaA), *dnaN* (encoding the beta subunit of DNA polymerase III) and *recF* (encoding the DNA repair protein RecF) or at least one of these three genes, are found directly upstream of *gyrB* gene in all bacteria in which *gyrB* is located in the immediate vicinity of *oriC*, such as most bacteria of Groups 1 and 3 (Table 1). Transcription from these co-oriented neighbouring genes provides a strong input of DNA relaxation (Sobetzko, 2016) that stimulates transcription of supercoiling-sensitive P_*gyrB*_ (Menzel & Gellert, 1987). This is true of most bacteria where *gyrB* is in the immediate vicinity of *oriC* and *gyrA* is located either about 20% of the chromosome further away or is a part of a *gyrBA* operon. Bacteria with a *gyrBA* operon that is not in the immediate vicinity of *oriC* (such as *Chlamydia psittaci*) and bacteria with the two *gyr* genes separated by about 20% of the chromosome, such as *Myxococcus xanthus* (together with some Bacteroidetes) that satisfy the gene positional parameters characteristic of Group 1, possess *gyrB* with a non-conserved genetic neighbourhood. Bacteria of Groups 2 and 4 do not have a conserved genetic environment around *gyrB*. The conservation of the genetic neighbourhood 5’ to *gyrB* seems to be more important than the subjective proximity to *oriC*. Therefore, genetic neighbourhood conservation was used as a parameter to decide the groupings in Table 1. The frequent association of *gyrB* with the *dnaA, dnaN* and *recF* genes; the higher conservation of *gyrB*’s position in comparison to *gyrA*; the transcriptional response of *gyrB* to quinolones and our inability to delete *gyrB* from the *gyrAB + gyrB* merodiploid *Salmonella* strain – all indicate that conservation of the physical location of *gyrB*, but not *gyrA*, is essential in many bacteria. These findings reveal important information about chromosome composition in natural bacteria and can help guide attempts at synthetic chromosome design.

### Experimental procedures

#### Bacterial strains and culture conditions

The bacterial strains used in this study were derivatives of *S*. Typhimurium strain SL1344 and their details are listed in Table 1. Bacteriophage P22 HT 105/1 *int-*201 was used for generalized transduction during strain construction (Schmieger, 1972). Phage lysates were filter-sterilized and stored at 4°C in the dark. Bacterial strains were stored as 35% glycerol stocks at −80°C and freshly streaked on agar plates for each biological replicate. Four ml LB broth was inoculated with a single colony and grown for 18 h. This overnight culture was sub-cultured into fresh 25 ml LB broth normalizing to an OD_600_ of 0.003, unless otherwise stated, and grown to the required growth phase. The standard growth conditions for all bacterial strains were 37°C, 200 rpm, unless otherwise stated. For culturing in minimal medium, overnight cultures were prepared as described above. 1 ml of overnight culture was washed three times with minimal medium N of the required MgCl_2_ concentration to remove nutrients, sub-cultured into minimal medium of the corresponding MgCl_2_ concentration in a total volume of 25 ml and grown for 24 h to pre-condition the bacteria. The pre-conditioned culture was sub-cultured into 25 ml of fresh minimal medium N adjusted to an OD_600_ of 0.03 and grown for a further 24 h to obtain a culture in the stationary phase of growth.

To measure growth characteristics of a bacterial culture, an overnight culture was adjusted to an OD_600_ of 0.003 in 25 ml of fresh LB broth and grown at the standard conditions for 24 h in the appropriate liquid medium. The optical density of the culture at OD_600_ was measured at 1-h intervals for the first 3 hours and then every 30 min until 8 hours; the last reading was taken at 24 h. Measurements were taken using a Thermo Scientific BioMate 3S spectrophotometer with liquid cultures in plastic cuvettes. To measure the growth characteristics of a bacterial culture in minimal medium with altered Mg^2+^ concentration, an overnight bacterial culture was washed in minimal medium with an appropriate concentration of MgCl_2_ and pre-conditioned for 24 h. The pre-conditioned culture was adjusted to an OD_600_ of 0.03 in 25 ml of fresh medium in two flasks and the OD_600_ was measured every hour beginning from 2 h post time zero until 8 h. Separate cultures were set up similarly to measure OD_600_ every hour from 8 h until 15 h. In this way, the number of times each flask was opened and sampled was minimized to yield reliable and reproducible measurements.

The growth characteristics of bacterial cultures in LB broth were also measured by viable counts. The culture was grown in the same way as for spectrophotometry, and an aliquot was taken at 2 h, 4 h, 6 h, 8 h and 24 h, serially diluted and spread on LB agar plates to give between 30 and 300 colonies after overnight incubation at 37°C. The bacterial colony counts were expressed as colony forming units per millilitre (cfu ml^-1^).

#### Bacterial motility assays

Assays were carried out precisely as described to achieve agreement between biological replicates. 0.3% LB agar was melted in a 100 ml bottle in a Tyndall steamer for 50 min, allowed to cool in a 55°C water bath for 20 min, six plates were poured and left to dry near a Bunsen flame for 25 min. 1 µl of bacterial overnight culture was pipetted under the agar surface with two inocula per plate. Plates were placed in a 37°C incubator without stacking to ensure equal oxygen access. After 5 h, the diameters of the resulting swarm zones were measured and expressed as the ratio of the WT zone to that of the mutant.

#### Competitive fitness assays

Flasks of broth were inoculated with the pair of competing bacterial strains in a 1:1 ratio. Derivatives of each competitor were constructed that carried a chloramphenicol acetyl transferase (*cat*) gene cassette within the transcriptionally silent pseudogene *SL1483*. This *cat* insertion is known to be neutral in its effects on bacterial fitness (Lacharme-Lora et al., 2019) and allows the marked strain to be distinguished from its unmarked competitor. Competitions were run in which wild type SL1344 was the marked strain or in which SL1344 *gyrBA* was the marked strain. Strains to be competed were pre-conditioned in separate 25 ml cultures for 24 h without antibiotics. Then 10^5^ cells of each strain were mixed in 25 ml of fresh LB broth and grown as a mixed culture for another 24 h. The number of colony forming units was determined by plating the mixture on chloramphenicol-containing plates and on plates with no antibiotic at T=0 h and T=24 h. Taking the wild type SL1344 Vs SL1344 *gyrBA* competition as an example, SL1344 was competed against SL1344 *gyrBA SL1483::cat* and, as a control, SL1344_*SL1483::cat* was competed against SL1344 *gyrBA*. The competitive fitness index (f.i.) was calculated according to the formula:

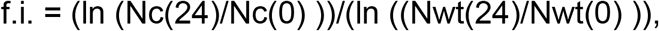

where Nc(0) and Nc(24) are the initial and final counts of a competitor and Nwt(0) and Nwt(24) are initial and final counts of the WT. Competitor is a strain other than the WT; f.i. <1 means that the competitor is less fit than the WT; f.i.>1 indicates the opposite.

#### *Construction of a* gyrBA *operon strain and an attempt to construct a* gyrAB *operon strain*

A derivative of *S*. Typhimurium with an artificial *gyrBA* operon was constructed by Lambda-Red homologous recombination (Datsenko & Wanner, 2000). Briefly, a *kan* cassette was amplified from plasmid pKD4 with primers Kan ins gyrA F and Kan ins gyrA R, (Table S1) using Phusion high-fidelity DNA polymerase. The amplicon, with overhangs homologous to a region immediately downstream of *gyrA*, was transformed into the WT strain harbouring plasmid pKD46, then grown in the presence of arabinose to activate the Lambda Red system in order to tag *gyrA* with the *kan* cassette. The *gyrA*::*kan* construct, including 20 nucleotides upstream from the *gyrA* translational start codon, was amplified using primers gyrB.int.gyrA::kan_Pf and gyrB.int.gyrA::kan_Prev (Table S1). The amplicon had overhangs that were homologous to sequences immediately downstream of *gyrB*. This allowed translation of the GyrA protein from the bicistronic *gyrBA* mRNA because several sequences closely matching to a consensus ribosome binding site (5’-AGGAGG-3’) were located in this 20 bp region. The *gyrA*::*kan* amplicon was inserted by Lambda Red-mediated recombination immediately downstream of the *gyrB* protein-coding region to construct the *gyrBA* operon. The original *gyrA* gene was deleted by an in-frame insertion of a *kan* cassette (Baba *et al*., 2006). The *kan* resistance cassettes were subsequently eliminated via FLP-mediated site-specific recombination (Cherepanov & Wackernagel, 1995). The resulting *gyrBA* strain had the genes that encode both subunits of DNA gyrase arranged as a bicistronic operon under the control of a common promoter, P_*gyrB*_ (Table 2).

**Table 2.**
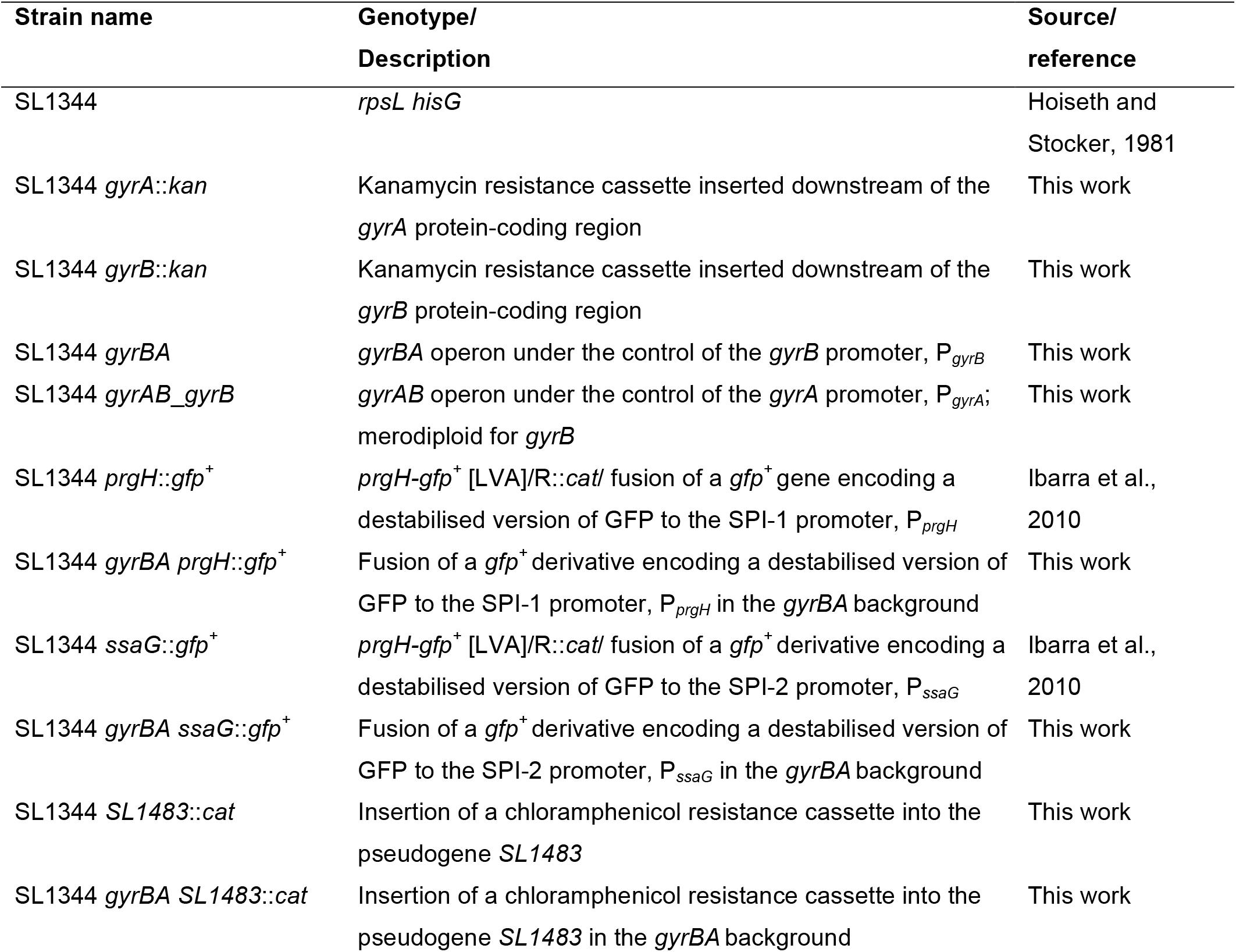
Bacterial strains

| Strain name | Genotype/<br>Description | Source/<br>reference |
| --- | --- | --- |
| SL1344 | *rpsL hisG* | Hoiseth and<br>Stocker, 1981 |
| SL1344 *gyrA::kan* | Kanamycin resistance cassette inserted downstream of the *gyrA* protein-coding region | This work |
| SL1344 *gyrB::kan* | Kanamycin resistance cassette inserted downstream of the *gyrB* protein-coding region | This work |
| SL1344 *gyrBA* | *gyrBA* operon under the control of the *gyrB* promoter, P<sub>*gyrB*</sub> | This work |
| SL1344 *gyrAB_gyrB* | *gyrAB* operon under the control of the *gyrA* promoter, P<sub>*gyrA*</sub>; merodiploid for *gyrB* | This work |
| SL1344 *prgH::gfp*<sup>+</sup> | *prgH-gfp*<sup>+</sup> [LVA]/R::*cat*/ fusion of a *gfp*<sup>+</sup> gene encoding a destabilised version of GFP to the SPI-1 promoter, P<sub>*prgH*</sub> | Ibarra et al.,<br>2010 |
| SL1344 *gyrBA prgH::gfp*<sup>+</sup> | Fusion of a *gfp*<sup>+</sup> derivative encoding a destabilised version of GFP to the SPI-1 promoter, P<sub>*prgH*</sub> in the *gyrBA* background | This work |
| SL1344 *ssaG::gfp*<sup>+</sup> | *prgH-gfp*<sup>+</sup> [LVA]/R::*cat*/ fusion of a *gfp*<sup>+</sup> derivative encoding a destabilised version of GFP to the SPI-2 promoter, P<sub>*ssaG*</sub> | Ibarra et al.,<br>2010 |
| SL1344 *gyrBA ssaG::gfp*<sup>+</sup> | Fusion of a *gfp*<sup>+</sup> derivative encoding a destabilised version of GFP to the SPI-2 promoter, P<sub>*ssaG*</sub> in the *gyrBA* background | This work |
| SL1344 *SL1483::cat* | Insertion of a chloramphenicol resistance cassette into the pseudogene *SL1483* | This work |
| SL1344 *gyrBA SL1483::cat* | Insertion of a chloramphenicol resistance cassette into the pseudogene *SL1483* in the *gyrBA* background | This work |

**Table 3.**
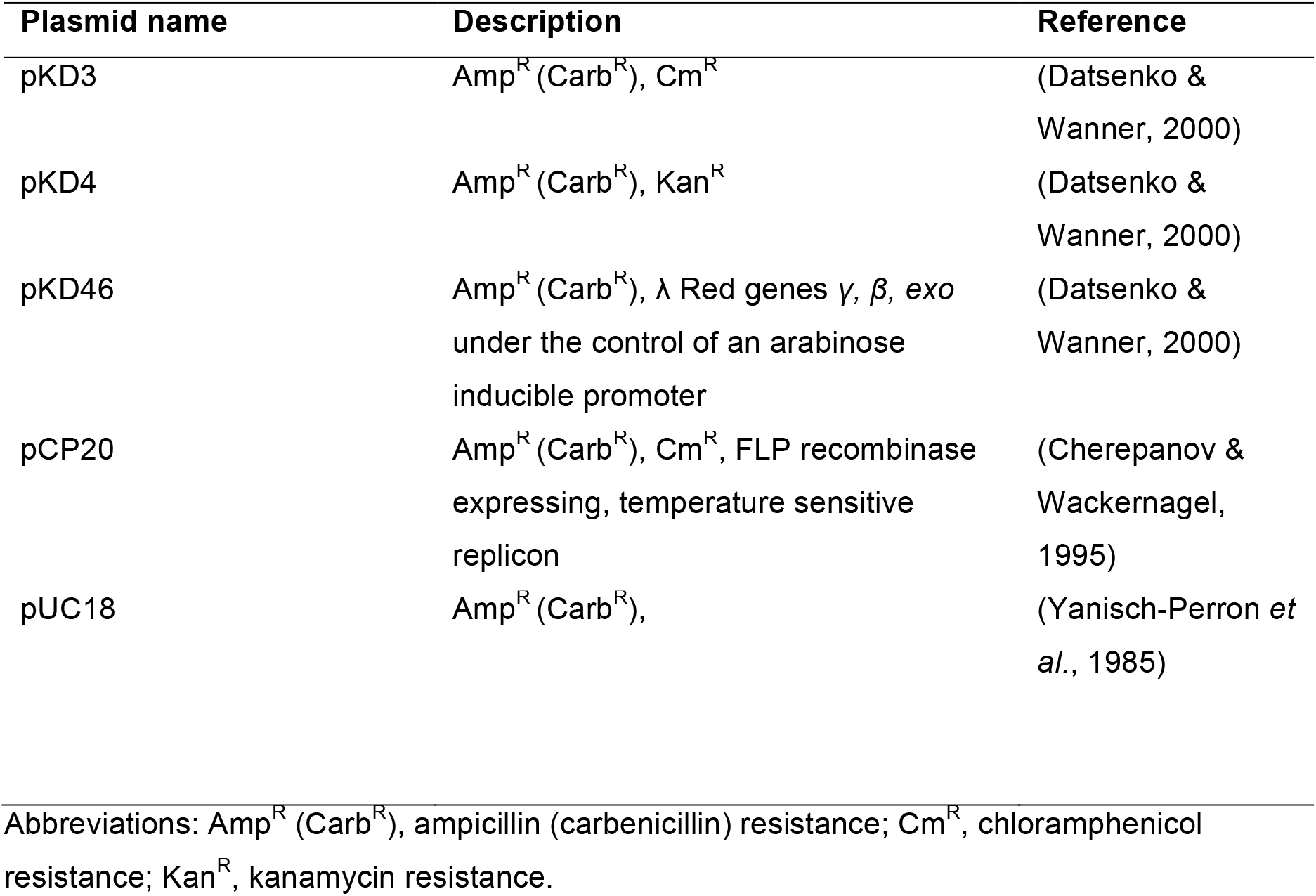
Plasmids used in this study

| Plasmid name | Description | Reference |
| --- | --- | --- |
| pKD3 | Amp<sup>R</sup> (Carb<sup>R</sup>), Cm<sup>R</sup> | (Datsenko & Wanner, 2000) |
| pKD4 | Amp<sup>R</sup> (Carb<sup>R</sup>), Kan<sup>R</sup> | (Datsenko & Wanner, 2000) |
| pKD46 | Amp<sup>R</sup> (Carb<sup>R</sup>), $\lambda$ Red genes $\gamma$, $\beta$, *exo* under the control of an arabinose inducible promoter | (Datsenko & Wanner, 2000) |
| pCP20 | Amp<sup>R</sup> (Carb<sup>R</sup>), Cm<sup>R</sup>, FLP recombinase expressing, temperature sensitive replicon | (Cherepanov & Wackernagel, 1995) |
| pUC18 | Amp<sup>R</sup> (Carb<sup>R</sup>), | (Yanisch-Perron *et al.*, 1985) |

Abbreviations: Amp<sup>R</sup> (Carb<sup>R</sup>), ampicillin (carbenicillin) resistance; Cm<sup>R</sup>, chloramphenicol resistance; Kan<sup>R</sup>, kanamycin resistance.

Although no *gyrAB* operon has been reported to occur naturally in bacteria, the construction of an artificial *gyrAB* operon in *Salmonella* was attempted in parallel with the *gyrBA* construction, using an identical strategy. The *gyrB* gene was tagged with a *kan* cassette that was amplified using primers Kan ins gyrB F and Kan ins gyrB R (Table S1). The *gyrB*::*kan* construct was amplified using primers gyrA.int.gyrB::Kan_Pf and gyrA.int.gyrB::Kan_Prev (Table S1) and inserted immediately downstream of *gyrA* to construct a *gyrAB* operon. To enable translation of the GyrB protein from the bicistronic *gyrAB* mRNA, 20 nucleotides upstream from the *gyrB* translation start codon were included in the insertion, providing *gyrA* with the best match to a consensus ribosome binding site found there: 5’-ACGAGG-3’. The resulting strain had a *gyrAB* operon at the *gyrA* locus and was merodiploid for *gyrB* (Table 2). Repeated attempts to delete *gyrB* from its original locus were unsuccessful. This indicated that the original position of *gyrB* is important and that it cannot be deleted even when *gyrB* is expressed from a synthetic operon located elsewhere on the chromosome. No further work was performed with the *gyrAB* operon strain.

#### DNA isolation for whole genome sequencing

To obtain high-quality chromosomal DNA for whole genome sequencing, a basic phenol-chloroform method was used (Sambrook & Russell, 2006). 2 ml of an overnight culture were centrifuged at 16000 x g for 1 min to harvest cells and the cell pellet was resuspended in 400 µl of TE buffer pH 8 (100 mM Tris-HCl pH 8.0, 10 mM EDTA pH 8.0 (BDH, Poole, England)). 1% SDS and 2 mg/ml proteinase K were added and incubated for 2 h at 37°C to complete lysis. DNA was isolated by the addition of 1 volume of phenol pH 8.0 : chloroform : isoamyl alcohol (25 : 24 : 1) (AppliChem, Darmstadt, Germany), thorough mixing and centrifugation at 16000 x g for 15 min at 4°C in phase-lock tube. The upper aqueous layer containing DNA was collected and the phenol: chloroform extraction was repeated two more times. To remove contaminants and to precipitate DNA, sodium acetate pH 5.2 at 0.3 M and isopropanol at 60% of the final volume were added and kept for 1 h at −20°C. DNA was pelleted by centrifugation at 16000 x g for 15 min at 4°C. The DNA pellet was washed with 70% ethanol, dried at 37°C until translucent and resuspended in 100 µl TE pH 8.0. The sample was electrophoresed on agarose gel to check for degradation and the DNA concentration was determined as follows: to remove RNA contamination from DNA samples, 100 mg/ml RNase A was added and incubated for 30 min at 37°C. Phenol-chloroform extraction was performed as above. To precipitate DNA, 0.5 M of ammonium acetate (Merck, Darmstadt, Germany) and a half volume of isopropanol were added and incubated for 2 h at −20°C. DNA was pelleted by centrifugation at 16000 x g for 15 min at 4°C. The DNA pellet was washed twice with 70% ethanol, dried at 37°C until translucent and resuspended in 50 µl water. The sample was run on an agarose gel to check for degradation. The concentration of DNA extracted was determined by measuring absorbance at 260 nm on a DeNovix DS-11 spectrophotometer (Wilmington, Delaware, USA). The shape of the absorbance curve was ensured to have a clear peak at 260 nm. The purity of samples was assessed by the ratio of A_260_/A_280_ – a measure of protein and phenol contamination and A_260_/A_230_ – a measure of contaminants such as EDTA, where both should be as close as possible to 2. Only high-quality samples were chosen for further work.

#### Whole genome sequencing

Whole genome sequencing was performed on final versions of the constructed strains to ensure that no compensatory mutations were introduced into their genomes. The sequencing was performed by MicrobesNG (Birmingham, UK) using Illumina next generation sequencing technology. The output reads were assembled using Velvet (Zerbino, 2010) and aligned to the reference SL1344 sequence NC_016810.1 Breseq software (Deatherage & Barrick, 2014). The data are available through the Sequence Read Archive (SRA) with accession number PRJNA682874.

#### RNA extraction, DNase treatment and RT-qPCR

RNA for measuring gene expression by qPCR was isolated using an acidic phenol-chloroform method. An overnight culture was subcultured into 25 ml of fresh LB broth normalising to an OD600 of 0.003. The bacterial culture was grown to the required timepoint and mixed with 40% volume of 5% acidic phenol (pH 4.3) in ethanol and placed on ice for at least 30 min to stop transcription. The cells were harvested by centrifugation at 3220 x g for 10 min at 4°C and resuspended in 700 µl of TE buffer pH 8 containing 0.5 mg/ml lysozyme. 1% SDS and 0.1 mg/ml proteinase K were added and incubated for 20 min at 40°C to complete lysis. 1/10 volume of 3 M sodium acetate was added to precipitate RNA, 1 volume of 1:1 solution of acidic phenol and chloroform was added, mixed well on a vortex mixer and centrifuged at 16000 x g for 15 min at 4°C to extract RNA into aqueous phase. To precipitate RNA the aqueous layer was harvested, mixed with 1 volume of isopropanol and incubated at −20°C for 1 hour. RNA was harvested by centrifugation at 16000 x g for 15 min at 4°C. The RNA pellet was washed with 70% ethanol and dried at 37°C until translucent. The total RNA was dissolved in 50 µl DEPC-treated water and its concentration was determined using DeNovix DS-11 spectrophotometer.

For DNase-treatment, RNA was diluted to 20 µg in 80 µl, denatured at 65°C for 5 min and kept on ice. 1x DNase I buffer including MgCl2 and 10 U DNase I (ThermoFisher Scientific, Waltham, US) were added and incubated for 45 min at 37°C. 100 µl of 1:1 acidic phenol : chloroform was added to DNase I digestion samples, mixed and transferred to a phase-lock tube. RNA was extracted by centrifugation at 16000 x g for 12 min at 15°C. The upper aqueous layer was harvested and RNA was precipitated by adding 2.5 volumes of 30:1 ethanol : 3 M sodium acetate pH 6.5 for 2 h or overnight at − 20°C. RNA was harvested by centrifugation at 16000 x g for 30 min at 4°C. The RNA pellet was washed with 70% ethanol and dried at 37°C until translucent. The total RNA was dissolved in 30 µl DEPC-treated water and its concentration was determined as in 2.5.5. RNA was checked for DNA contamination by the end point PCR and for integrity on a HT gel (Mansour & Pestov, 2013).

400 nm of the total extracted and DNase I treated RNA was converted to cDNA using GoScriptTM Reverse Transcription System kit (Promega) according to manufacturer’s guidelines. Then, 5.33 ng of cDNA in 20 µl reaction was used as a template for Real Time quantitative PCR (RT-qPCR) using 1x FastStart Universal SYBR Green Master (ROX) (Roche, Mannheim, Germany) and gene-specific pair of primers (0.3 µM each). For each pair of primers, a standard curve was generated using 10-fold serially diluted gDNA. PCR and fluorescence detection were carried out in StepOne Real Time PCR system (Applied Biosystems). Analysis was performed in the accompanying software. The cycling conditions were as follows: 10 min at 95°C; 40 cycles of 15 sec at 95°C and 1 min at 60°C.

#### Minimum inhibitory concentration (MIC) of antibiotics determination

MIC90 of antibiotics (a minimal concentration at which 90% of bacterial growth is inhibited) was found by serially diluting antibiotics and spectrophotometrically testing the ability of different dilutions to inhibit bacterial growth. On a 96-well plate, all wells (excluding column 12) were filled with 60 µl of sterile LB broth. 1 ml solutions of antibiotics to be tested were prepared at the highest desired concentration in LB. 300 µl of the prepared antibiotics were added to the wells of column 12 and homogenised by pipetting up and down 5 times with a multichannel pipette. 240 µl was transferred to the next wells in column 11, homogenisation was repeated and serial 1:1.25 dilutions were sequentially continued until column 3. The final 240 µl from column 3 were discarded. All the wells were inoculated with bacterial cultures adjusted to an OD_600_ of 0.003 except column 1. In this way column 1 contained negative controls (no bacteria and no antibiotic), column 2 contained positive controls (no antibiotics) and columns 3-12 contained serially diluted antibiotics inoculated with the identical number of bacteria. The plate was covered, sealed between plastic sheets and incubated for 18 h at the standard growth conditions. The plate was read by measuring OD_600_ values on a plate reader (Multiscan EX, Thermo Electronics).

#### SPI-1 and SPI-2 reporter assays

*Salmonella* pathogenicity island (SPI) activity was accessed by measuring the expression of *gfp*^*+*^ reporter gene fusions to promoters of *prgH* and *ssaG* to look at SPI-1 and SPI-2 expression, respectively. The *gfp*^*+*^ reporter fusions were transduced into each strain by P22 generalized transduction and selected with chloramphenicol. 100 µl of overnight culture of the *gfp*^*+*^ reporter-carrying strain was diluted 1:100 in LB broth. Black 96 plate with transparent flat bottom was filled with 100 µl of the diluted culture in six technical replicates, negative controls were included. The plate was sealed with parafilm and incubated at 300 rpm, 37°C for 24 h in the Synergy H1 microplate reader (Biotek, Vermont, USA). Bacterial growth was measured at 600 nm and GFP fluorescence was read using 485.5 nm excitation frequency at 528 nm emission frequency, measurements were taken every 20 min. For measurements in the minimal medium, the culture was adjusted to OD_600_ of 0.03 in the medium of the required MgCl_2_ concentration and measurements proceeded as above.

#### Global supercoiling determination

Global DNA supercoiling was assayed in bacterial strains transformed with a reporter plasmid pUC18. An overnight culture of pUC18-containg strain was adjusted to an OD_600_ of 0.003 and grown to the late stationary growth stage (24 h) in 25 ml LB broth or in 25 ml of minimal medium N of the required MgCl_2_ concentration pre-conditioned as above. Fourteen OD_600_ units (6 OD_600_ units for minimal medium) were harvested and pUC18 was isolated with the aid the of QIAprep Spin miniprep kit (QIAGEN, Hilden, Germany) according to manufacturer’s guidelines.

To observe the range of DNA supercoiling states characteristic of a strain at a given growth stage, extracted pUC18 samples were resolved on 0.8% agarose gel supplemented with the DNA intercalating agent chloroquine. 2 L of 1x TBE buffer (89 mM Tris base, 89 mM boric acid, 2 mM EDTA pH 8.0) and 1 ml of 25 mg/ml chloroquine were made. 0.8% agarose solution was made from 300 µl TBE and melted in a Tyndall steamer. When the gel cooled down, it was supplemented with 2.5 µg/ml chloroquine. The 27 cm long gel was poured, left to polymerise for 2 h and covered with 1.7 litres of the running buffer containing 1x TBE and chloroquine at 2.5 µg/ml. 1 µg or 500 ng of the plasmid samples in 15 µl volumes were mixed with 5x loading dye (80% glycerol, 0.01% bromophenol blue) and loaded on a gel. The gel was electrophoresed for 16 h at 100 V. The gel was washed in distilled water for 24 h changing water a few times, stained in 1 µg/ml ethidium bromide for 1 h rocking in the dark. The stain was poured off and the gel was washed in distilled water for further 1 h. The plasmid topoisomers were visualised under UV on the ImageQuant LAS 4000 imager. ImageJ software was used to outline plasmid topoisomer distribution profiles.

#### *Determining the patterns of* gyrA *and* gyrB *locations in bacterial chromosomes*

The location of *oriC* in each organism examined was determined using the DoriC 10.0 database (tubic.org/doric) (Luo and Gao, 2019) and the coordinates of their *gyrA* and *gyrB* genes were obtained using the Ensembl bacteria browser (bacteria.ensembl.org). Distance in base pairs between the *oriC* and the gene was calculated and converted into the percentage of the total chromosome size. An attempt was made to cover bacterial taxonomy as broadly as possible, encompassing members of the major bacterial phyla, well studied, and clinically important organisms in the analysis (Table 1). The table is neither complete nor does it claim to include all the existing possibilities of *gyrA* and *gyrB* arrangements in bacterial chromosomes, but instead, exemplifies the arrangement possibilities mentioned in this work. Closely related species and those belonging to the less diverse phyla were found to share the chromosomal positions of *gyrA* and *gyrB* frequently. Thus, one representative of a taxonomic rank was often deemed sufficient for the purpose of inclusion in the table. Lower classification ranks were analysed within more diverse and studied phyla.

#### Mammalian cell culture conditions

RAW264.7 murine macrophages were maintained in Dulbecco’s Modified Eagle’s Medium (DMEM), (Sigma, catalogue number D6429) supplemented with 10% fetal bovine serum (FBS) in a humidified 37°C, 5% CO_2_ tissue-culture incubator grown in 75 cm^3^ tissue-culture flasks. When approximately 80% confluent growth was achieved, cells were split to a fresh flask. Cells within the 9-16 passage number range were used for infections. All media and PBS used for cell culture were pre-warmed to 37°C. To split cells, old DMEM was removed and the monolayer was rinsed with 10 ml of sterile PBS. Ten ml of fresh DMEM was pipetted into the flask and the monolayer was scraped gently with a cell scraper to dislodge the cells. Scraped cells were centrifuged at 450 x g for 5 min in an Eppendorf 5810R centrifuge and the cell pellet was resuspended in 5 ml DMEM+FBS. One ml of the cell suspension was added to 14 ml of fresh DMEM+FBS in a 75 cm^3^ flask, gently rocked to mix and incubated at 37°C, 5% CO_2_. To seed cells for infection, cells were treated as for splitting. After resuspension in 5 ml DMEM+FBS, viable cells were counted on a haemocytometer using trypan blue exclusion dye. A 24-well tissue culture plate was filled with 500 µl DMEM+FBS. 1.5×10^5^ cells were added to each well, gently rocked to mix and incubated at 37°C, 5% CO_2_ for 24 h.

#### Macrophage viability assay in SPI-1 inducing conditions

Overnight bacterial cultures were subcultured 1:33 in 10 ml of fresh LB broth in 125 ml conical flask and grown for 3.5 h to maximize SPI-1 expression (Steele-Mortimer *et al*., 1999). 500 µl of the culture was centrifuged at 16000 x g for 1 min and resuspended in 500 µl of HBSS-/-. Monolayers were washed twice with 500 µl of HBSS+/+ and infected with bacteria at MOI of 5 in three technical replicates for each timepoint and strain. The plate was centrifuged at 200 x g for 10 min to synchronize infections and incubated for 30 min at 37°C, 5% CO_2_. In the meantime, the infection medium was plated for enumeration on LB agar plates – T=0 h. Gentamycin protection assay was used to determine bacterial counts inside macrophages. To kill all extracellular bacteria, the monolayers were washed once with HBSS+/+ and high gentamycin (100 µg/ml) treatment diluted in DMEM+FBS was added to the wells. The plate was incubated at 37°C, 5% CO_2_ for 1 h. At 1 h post infection the monolayers were washed three times with HBSS+/+, macrophages were lysed by adding 1 ml of ice-cold water, pipetting up and down ten times with scraping and intracellular bacteria were plated for enumeration. The monolayers which were intended for other timepoints, were washed once with HBSS+/+, low gentamycin (10 µg/ml) treatment in DMEM+FBS was added and the plate was incubated at 37°C, 5% CO_2_. The low gentamycin concentration is to ensure that any extracellular bacteria are killed, but at the same time to avoid gentamycin permeabilizing plasma membrane of a macrophage (Kaneko *et al*., 2016). At later timepoints monolayers were washed three times with HBSS+/+, macrophages were lysed by adding 1 ml ice-cold water, pipetting up and down ten times with scraping and intracellular bacteria were plated for enumeration.

## Acknowledgements

Research in the CJD laboratory is supported by Science Foundation Ireland Investigator Award 13/IA/1875.

## SUPPLEMENTARY FILES

**Fig. S1.**
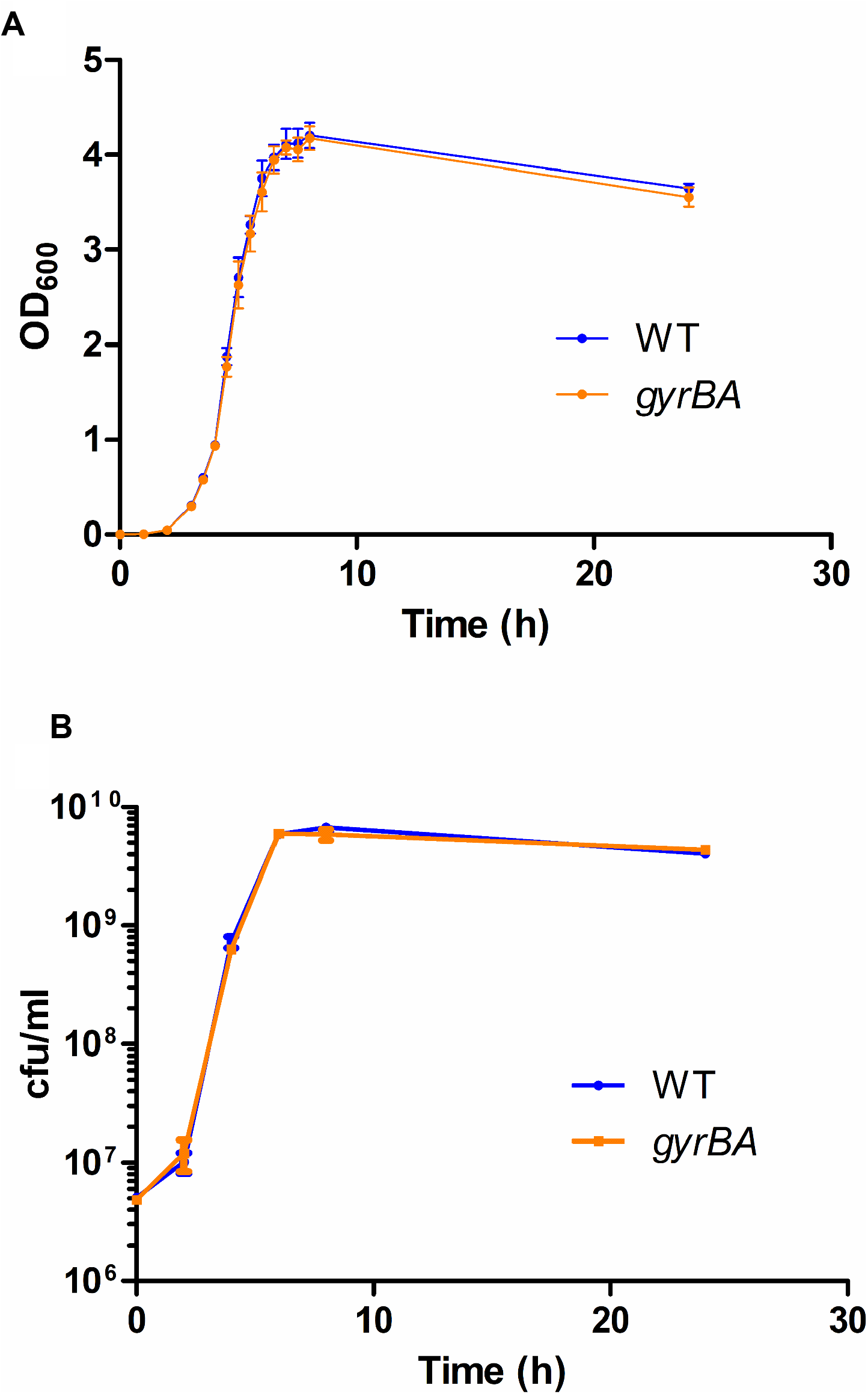
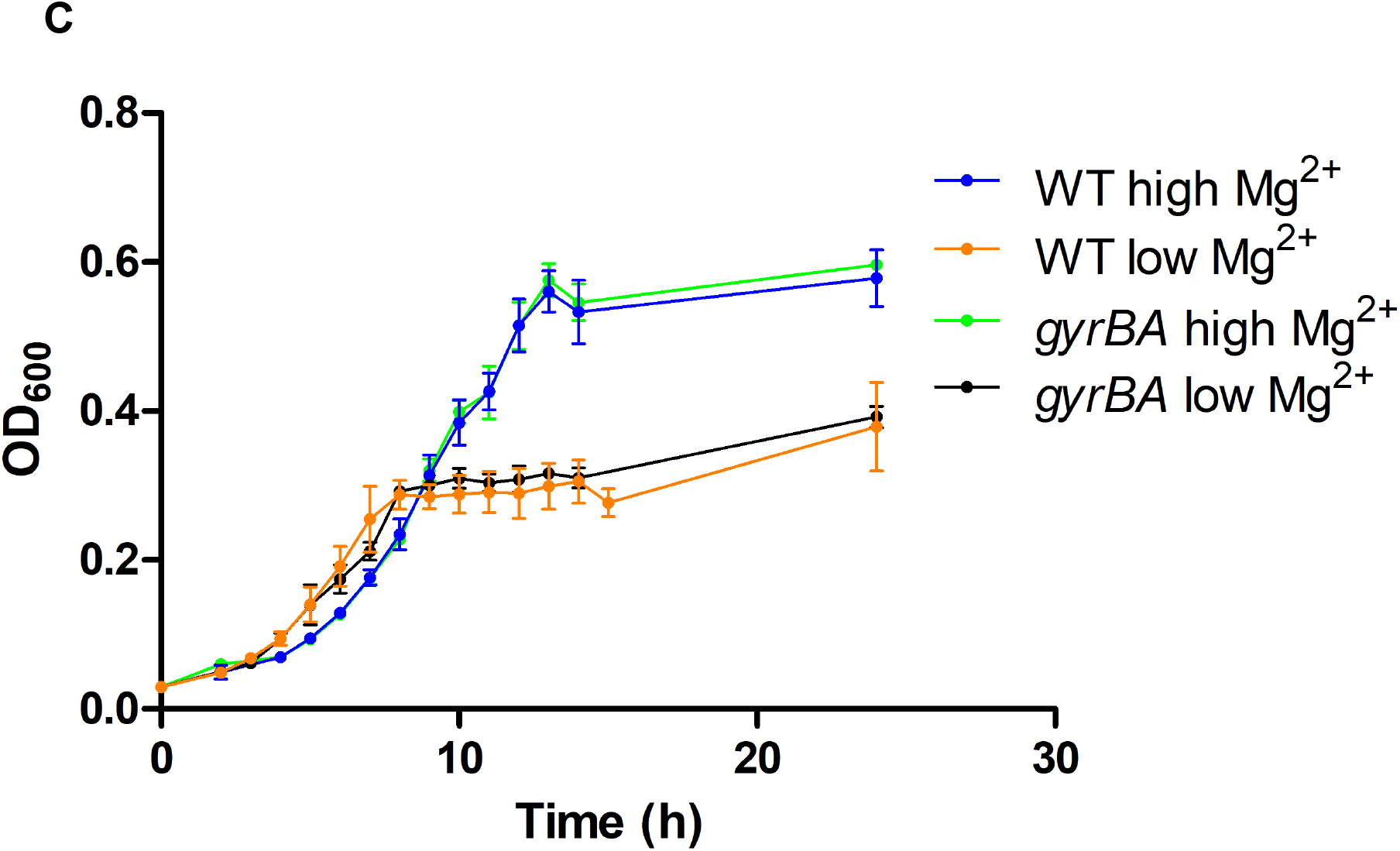
Growth characteristics of SL1344 *gyrBA*. A. Growth of the *gyrBA* strain as measured by absorbance at 600nm. OD_600_ measurements were made every hour until 3 h, then every 30 min until 8 h and lastly at 24 h. B. Growth of the *gyrBA* strain as measured by viability counts. Dilutions of bacterial cultures were spread on agar plates, incubated at 37°C and colonies were counted. C. Growth of the WT and the *gyrBA* strains as measured by absorbance at 600 nm in minimal medium N. Pre-conditioned culture was subcultured into 25 ml of fresh minimal medium of the required Mg^2+^ concentration, normalizing to an OD_600_ of 0.03. OD_600_ values were measured every hour from 2 h until 15 h and at 24 h. All plots are the results of at least three biological replicates, error bars represent standard deviation.

**Fig. S2.**
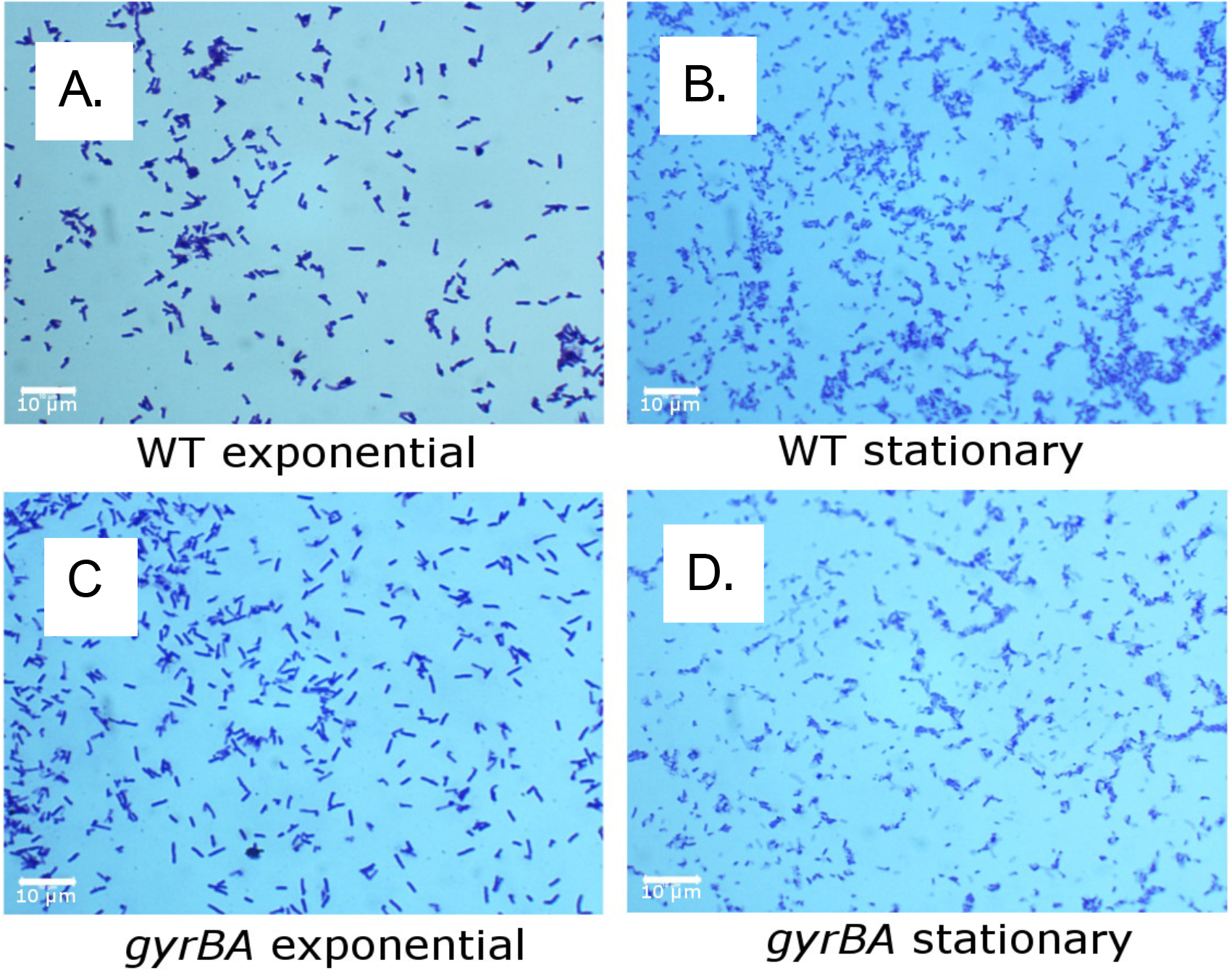
Cell morphology of SL1344 *gyrBA* at the exponential and stationary phases of growth. Bacteria were harvested at the mid-exponential or at the late stationary phases of growth, washed with PBS, heat-fixed, stained with crystal violet and viewed under 1000x magnification with an oil immersion lens. Standard rod-shaped *Salmonella* cells were observed in the WT and the *gyrBA* strains. All images are representative of three biological replicates. A 10 µm scale bar is given for reference. Cell morphology in LB liquid cultures is shown for wild type SL1344 in exponential phase (A), stationary phase (B), SL1344 *gyrBA* in exponential phase (C) and stationary phase (D).

**Fig. S3.**
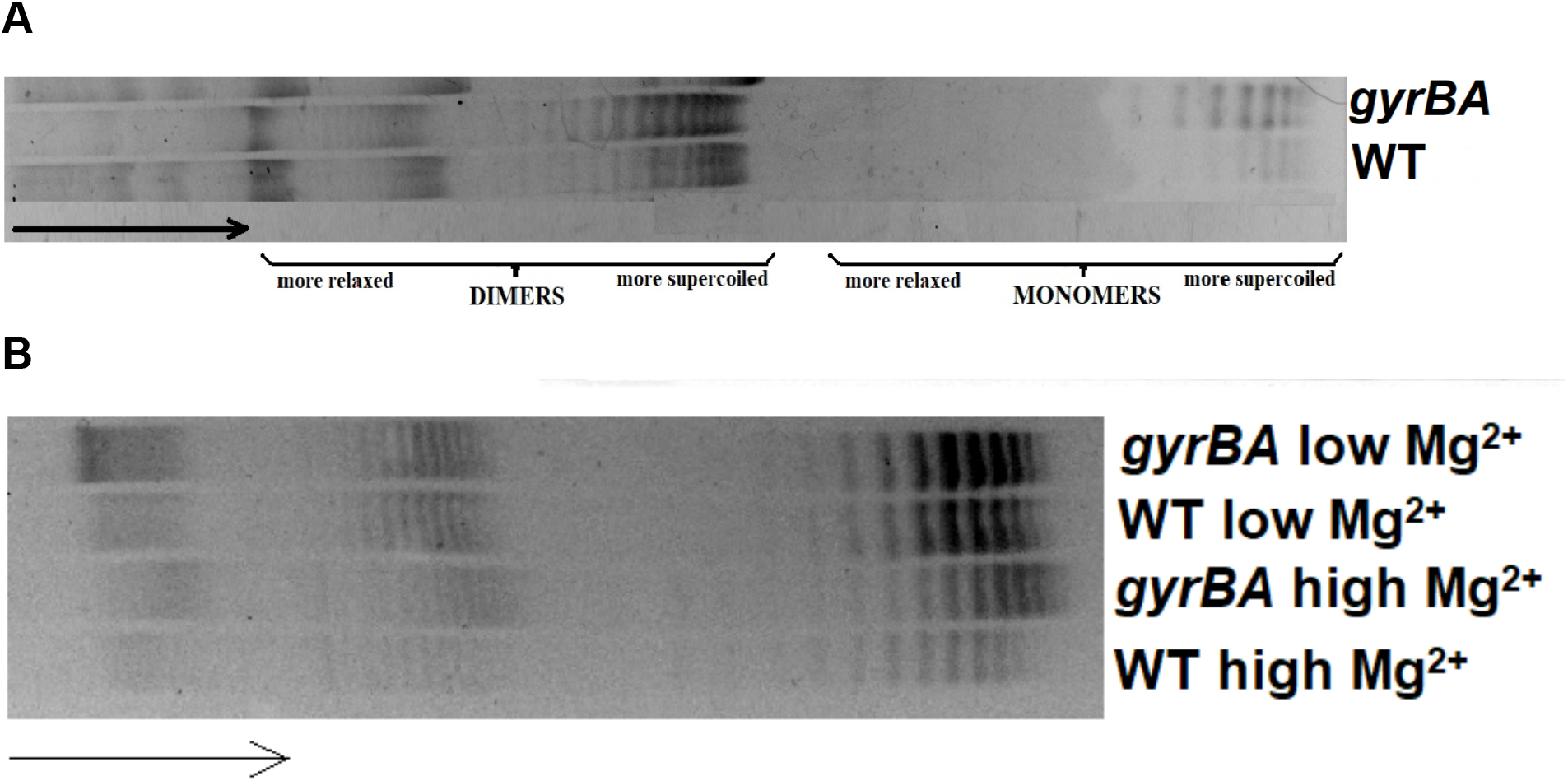
Full gel data for reporter plasmid DNA supercoiling in SL1344 and SL1344 *gyrBA*. Samples of pUC18 plasmid were extracted from the WT and the *gyrBA* at the stationary growth phase and run on a 0.8% agarose gel containing 2.5 µg/ml chloroquine to separate pUC18 topoisomers according to the degree of DNA supercoiling. Arrow shows the direction of electrophoretic flow with the more supercoiled plasmid topoisomers at the right of the gel. A) Global DNA supercoiling pattern of the WT and the *gyrBA* when grown in LB. The positions of relaxed and supercoiled plasmid topoisomer dimers and monomers are shown. B) Global DNA supercoiling pattern of the WT and the *gyrBA* when grown in minimal medium N with high (10 mM) Mg^2+^ and low (10 µM) Mg^2+.^ Plasmid topoisomer monomers are at the righthand end of the distribution; dimers and higher order oligomers are to the left.

**Table S1.**
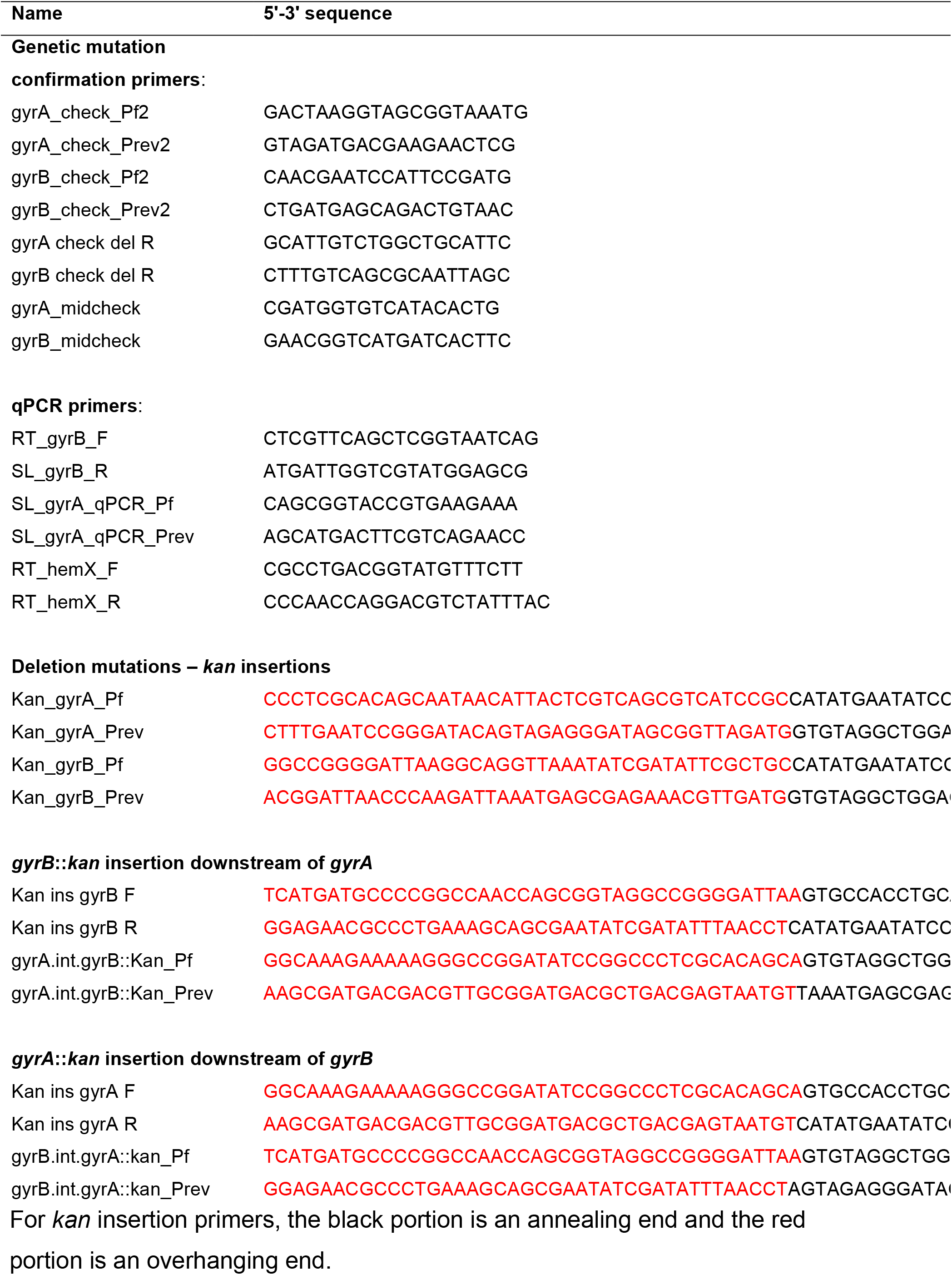
Oligonucleotides used in this study

